# Cryo-EM structures of the BAM–P1/P2-visible SurA complex reveal dynamic and cooperative interactions in outer membrane protein assembly

**DOI:** 10.1101/2025.11.10.687524

**Authors:** Ryoji Miyazaki, Hidetaka Kohga, Nami Matsuoka, Yuki Maruno, Wataru Yoshimoto, Yutaro S. Takahashi, Dede Heri Yuli Yanto, Yudhi Nugraha, Hideki Shigematsu, Takuya Shiota, Tomoya Tsukazaki

**Affiliations:** Nara Institute of Science and Technology, Ikoma, Nara, Japan; Interdisciplinary Graduate School of Agriculture and Engineering, University of Miyazaki, 1-1 Gakuen-Kibanadai Nishi, Miyazaki, Japan; Frontier Science Research Center, University of Miyazaki, 5200 Kihara, Kiyotake, Miyazaki, Japan; Research Center for Applied Microbiology, National Research and Innovation Agency (BRIN), Cibinong, Bogor, West Java, Indonesia; Research Center for Molecular Biology Eijkman, National Research and Innovation Agency (BRIN), Cibinong, Bogor, West Java, Indonesia; Structural Biology Division, Japan Synchrotron Radiation Research Institute, Sayo, Hyogo, Japan

## Abstract

The outer membrane (OM) of Gram-negative bacteria acts as a permeability barrier against toxic compounds. Its integrity is maintained by various outer membrane proteins (OMPs), which are inserted into the OM by the β-barrel assembly machinery (BAM) complex. The periplasmic chaperone SurA delivers unfolded OMPs to BAM; however, the mechanism of substrate transfer remains unclear. Here, we show that the flexible P1 and P2 domains of SurA regulate the function of its Core domain and interact with BAM components, including BamE, whose interaction with the P2 domain is crucial for efficient OMP assembly. Moreover, cryo-electron microscopy revealed four distinct *Escherichia coli* SurA–BAM structures, suggesting dynamic domain rearrangements of SurA. Based on these findings, we propose a dynamic model in which SurA transfers substrates to BAM through multiple conformational changes, providing a unified framework for chaperone-assisted OMP biogenesis.

## Introduction

Gram-negative (diderm) bacteria possess the inner membrane (IM) and the outer membrane (OM). The periplasmic space between the two membranes contains a mesh-like peptidoglycan cell wall that provides structural rigidity to the cell envelope. The OM serves as the interface with the external environment, functioning as a permeability barrier against toxic compounds, including antibiotics, and contributing to virulence and pathogenesis^1^. Lipopolysaccharides (LPSs) in the outer leaflet and densely packed β-barrel outer membrane proteins (OMPs) are essential for maintaining the integrity and function of the OM. OMPs are synthesized in the cytoplasm and translocated across the IM through the Sec translocon (**Supplementary Fig. 1a**)^2,3^. In the periplasm, they interact with periplasmic chaperones, such as SurA, Skp, FkpA, and PpiD, in an unfolded state and are subsequently delivered to the β-barrel assembly machinery (BAM) complex^4,5^. The BAM complex then mediates the folding and insertion of OMPs into the OM.

In *Escherichia coli*, the BAM complex consists of one β-barrel OMP, BamA, and four OM lipoproteins, BamB, BamC, BamD, and BamE^6,7^. BamA, the central component of the BAM complex, can catalyze OMP assembly. BamA consists of a 16-stranded transmembrane β-barrel domain at its C-terminus and a periplasmic region containing five polypeptide transport-associated (POTRA) domains at its N-terminus^6,7^. BamA and BamD are essential for cell viability and play critical roles in substrate recognition ^8–13^. Although their detailed roles remain unclear, the other nonessential Bam factors, BamB, BamC, and BamE, contribute to the function and stability of the BAM complex^14,15^. The BAM-mediated OMP assembly mechanism has been widely studied. Cryo-electron microscopy (Cryo-EM) studies have determined many structures of the BAM complex in apo states or bound to substrate proteins, inhibitors, and associated factors^6–12,16–19^. Based on these structural and biochemical analyses, a model for BAM-mediated OMP assembly has been proposed. After being transferred to the BAM complex, the C-terminal β-signal and internal signal of the OMP intermediate are first engaged by BamD, after which the β-signal is recognized by the N-terminal strand of the BamA β-barrel^8,10,12,13^. The nascent OMP β-barrel is then folded and inserted into the OM through the lateral gate of BamA. Upon release from the BAM complex, the substrate OMP completes β-barrel closure to achieve its mature conformation^11^. Although the molecular mechanism of BAM-mediated OMP assembly has been extensively characterized, it remains unclear how substrate OMPs are delivered to the BAM complex. Among the periplasmic chaperones involved in OMP biogenesis, SurA plays a major role in delivering unfolded OMPs to the BAM complex.

SurA is a key player in periplasmic chaperone networks. Deletion of *surA* causes many phenotypes, including reduced cellular accumulation of most OMPs, induction of stress responses, defects of OM integrity, and decreased pathogenicity^20–23^. Thus, SurA is considered to be the primary chaperone responsible for delivering unfolded OMP substrates to the BAM complex. In *E. coli*, SurA consists of three domains: a Core domain composed of the N- and C-terminal regions, and two parvulin-like peptidyl prolyl isomerase (PPIase) domains (P1 and P2 domains) (**Supplementary Fig. 1b**)^24^. Among these domains, only the Core domain is essential for SurA function *in vivo*^25^. The Core and P1 domains have been shown to interact with unfolded substrate, and SurA preferentially recognizes aromatic-enriched sequences within these substrates^26–30^. The flexible P1 and P2 domains undergo conformational changes that may modulate or support SurA function^31^. SurA also directly interacts with the BAM complex. Recent cryo-EM studies have determined several structures of the SurA–BAM complex (**Supplementary Fig. 1c**)^16,17^. In these structures, the Core domain of SurA interacts with the POTRA1 domain of BamA, which is important for BAM function. These studies also suggested that SurA can undergo conformational rearrangements upon binding to the BAM complex. Although the Core and P1 domains were resolved in these structures, the P2 domain was not. Thus, it remains unclear how SurA transfers its substrates to the BAM complex.

Here, we aimed to elucidate the molecular mechanism for SurA-mediated substrate transfer to the BAM complex. We found that the flexibly movable P1 and P2 domains of SurA regulate the activity of its Core domain. Further, we resolved four distinct structures of the SurA–BAM complex, including P2-visible and P1/P2 -visible structures, by cryo-EM. Crosslinking analyses identified SurA–BAM interaction sites and suggested that SurA interacts with the BAM complex as in our cryo-EM structures. We also show that the SurA–BAM interaction, especially the interaction between the SurA P2 domain and BamE, is important for OMP assembly. Based on these findings, we propose a dynamic model of substrate delivery in which conformational changes of SurA on the BAM complex mediate the transfer of substrate OMPs.

## Results

### Flexible P1/P2 regions of SurA regulate the function of the Core domain

To elucidate the molecular basis of SurA-mediated substrate delivery to the BAM complex, structural information on the BAM/SurA complex is essential. To stabilize the SurA–BAM interaction, we fused SurA to the N-terminus of the mature region of BamA via a Gly-Gly-Ser-Gly linker (**Supplementary Fig. 2a**). The fusion construct was coexpressed with BamB, BamC, BamD, and BamE-His8 in *E. coli* and purified as the SurA–BAM complex (**Supplementary Fig. 2b, c**). Cryo-EM single-particle analysis revealed two distinct structures of the SurA–BAM complex (**Fig. 1a**; **Supplementary Fig. 2d–f**). In one structure, only the Core domain of SurA was visible (referred to as the “Core-only” structure) (**Fig. 1a**, left), whereas in the other, the Core and P1 domains were resolved (referred to as the “P1-visible” structure) (**Fig. 1a**, right). Recently, two independent groups have reported highly similar structures (**Supplementary Fig. 3**). In both of our structures, the N-terminal region of the SurA Core domain interacts with the POTRA1 domain of BamA and contacts BamB. This SurA–BamA interaction is mediated via β-augmentation and is functionally important^16,17^. In the P1-visible structure, the P1 domain of SurA additionally contacts BamB. Thus, the P1 domain of SurA can adopt a distinct conformation that enables an additional interaction with BamB. The P2 domain was disordered in these structures. These observations suggest that the P1 and P2 domains of SurA are intrinsically flexible and undergo dynamic conformational changes.

**Fig. 1.**
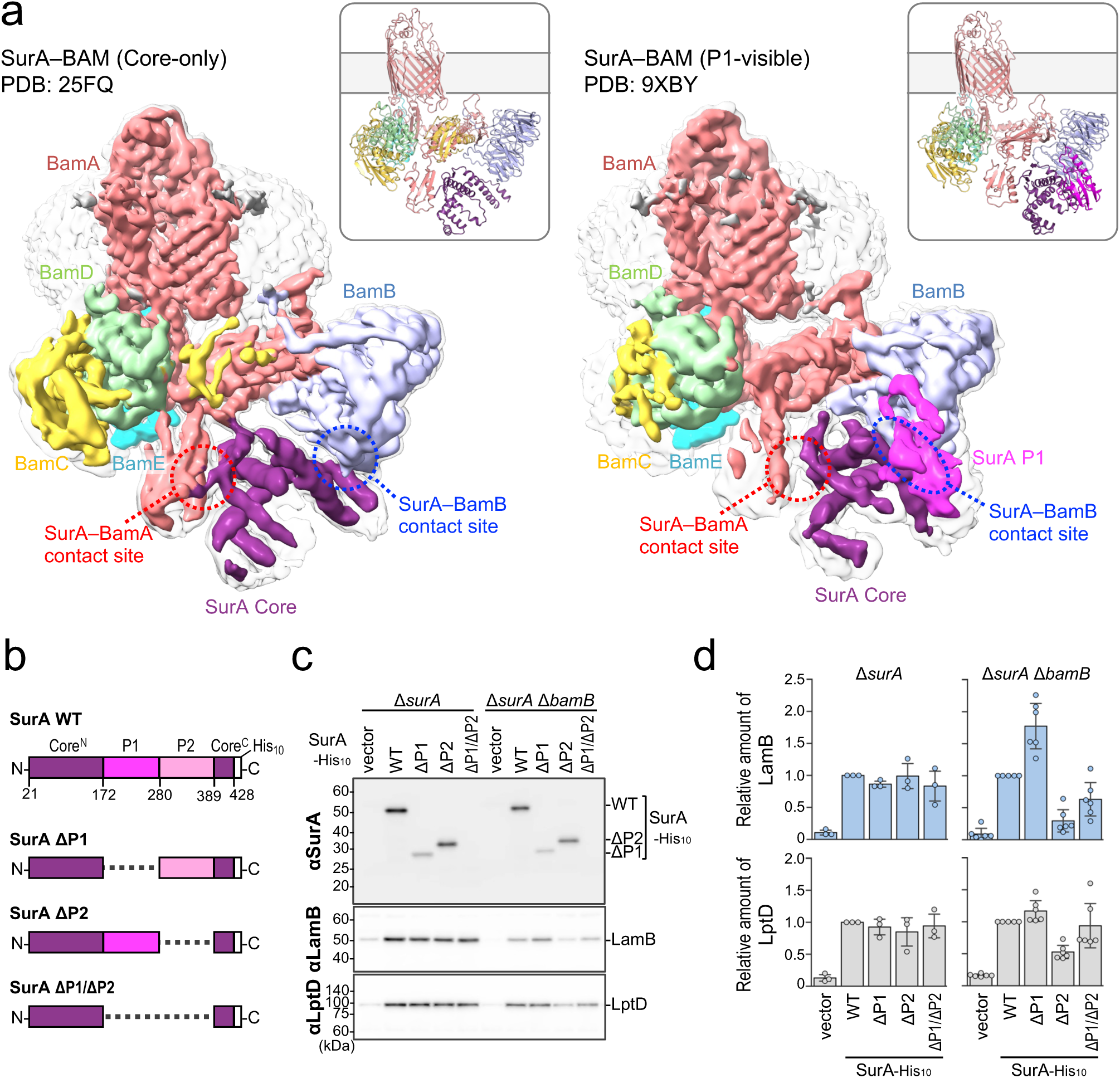
Flexible P1 and P2 regions of SurA regulate function of the Core domain. **a** Cryo-EM structures of SurA-BAM complex. Cryo-EM maps and cartoon models of SurA-BAM complexes are shown (Core-only, PDB: 9XD7; and P1-visible PDB: 9XBY). The BAM complex consists of BamA (red), BamB (light blue), BamC (yellow), BamD (light green), and BamE (cyan). The SurA Core (purple) is seen in both structures, and the P1 domain (magenta) only in the P1-visible one. Contact sites of SurA with BamA and BamB are shown in red and blue and circles, respectively. **b** Schematic representations of SurA deletion mutants lacking P1 and/or P2 domains. **c**, **d** Functional analysis of SurA deletion mutants. RM5457 (Δ*surA*) or RM5477 (Δ*surA*, Δ*bamB*) cells carrying pHM1021 (vector) or pHM1021-*surA(mut)-his10* plasmids were grown at 30 °C in LB medium supplemented with 1 mM IPTG for 2.5 h to express the indicated SurA-His10 variants. Total cellular proteins were acid-precipitated and analyzed by 10% Laemmli SDS‒PAGE followed by immunoblotting using the indicated antibodies. The amounts of LamB and LptD are quantified and shown relative to those of wild-type SurA-His10-expressing cells in (**d**) (n ≥ 3, ± S.D.).

To examine P1 and P2 functions, we constructed SurA mutants lacking P1 and/or P2 domains (ΔP1, ΔP2, and ΔP1/P2 mutants; **Fig. 1b**). In Δ*surA* cells, the cellular levels of the OMPs LamB and LptD were decreased (**Fig. 1c**, vector). Overexpression of wild-type (WT) SurA complemented this phenotype, as LamB and LptD accumulated normally in Δ*surA* cells (**Fig. 1c**, WT). The ΔP1, ΔP2, and ΔP1/P2 mutants also complemented the phenotype (**Fig. 1c, d**). These results indicate that the Core domain alone is sufficient for SurA function, consistent with previous findings^25^. As the Core and P1 domains of SurA physically interact with BamB, SurA may functionally cooperate with BamB. Therefore, we examined the effects of the SurA mutants in Δ*surA* Δ*bamB* cells. Interestingly, LamB accumulated at higher levels in cells expressing the ΔP1 mutant and at lower levels in the ΔP2-expressing cells compared with WT SurA (**Fig. 1c, d**). For LptD, the effects of the mutants were similar to those observed for LamB, albeit less pronounced. These results suggest that the P1 domain of SurA plays an inhibitory role, whereas the P2 domain plays a facilitatory role. Considering that the Core domain alone is essential for SurA function, the P1 and P2 domains likely regulate the activity of the Core domain.

### P1/P2 domains of SurA physically and functionally interact with Bam factors

The P1 domain of SurA contacts BamB in the P1-visible SurA–BAM structure (**Fig. 1a**), suggesting that the regulatory roles of the P1 and P2 domains are mediated through their interactions with the BAM complex. Therefore, we performed a systematic *in vivo* photo-crosslinking analysis to identify the BAM-contacting sites within SurA. We introduced *p*-benzoyl-L-phenylalanine (*p*BPA) at each of 41 positions (approximately every ten residues) in the mature region of SurA-His10 and carried out photo-crosslinking experiments (**Fig. 2a**). Δ*surA* cells expressing SurA(*p*BPA)-His10 were grown, UV-irradiated, and analyzed by immunoblotting with anti-SurA antibodies (**Supplementary Fig. 4**). Several SurA(*p*BPA) derivatives generated crosslinked products with higher apparent molecular masses than SurA-His10. Immunoblotting with anti-BamA antibodies revealed that SurA D26*p*BPA, K326*p*BPA, and N336*p*BPA derivatives were crosslinked to BamA (**Fig. 2b**). With anti-BamB antibodies, SurA E156*p*BPA, N166*p*BPA, A246*p*BPA, and E416*p*BPA derivatives were crosslinked to BamB (**Fig. 2c**). Furthermore, since a small crosslinked band observed for the SurA P296*p*BPA disappeared in the Δ*bamE* background (**Fig. 2d**), this band corresponds to a SurA–BamE crosslinked product. It should be noted that a band observed at around 100 kDa is likely to represent a nonspecific dimer of SurA. These results identified the crosslinking sites of SurA with the BAM complex (**Fig. 2a**) and revealed that the P1 and P2 domains of SurA interact not only with BamA, the central component of the BAM complex, but also with the accessory factors BamB and BamE. Mapping these residues onto our Core-only and P1-visible SurA–BAM structures showed that they are indeed located near BamA and BamB (**Fig. 2e**). Therefore, SurA appears to interact with the BAM complex in a manner consistent with these structures.

**Fig. 2.**
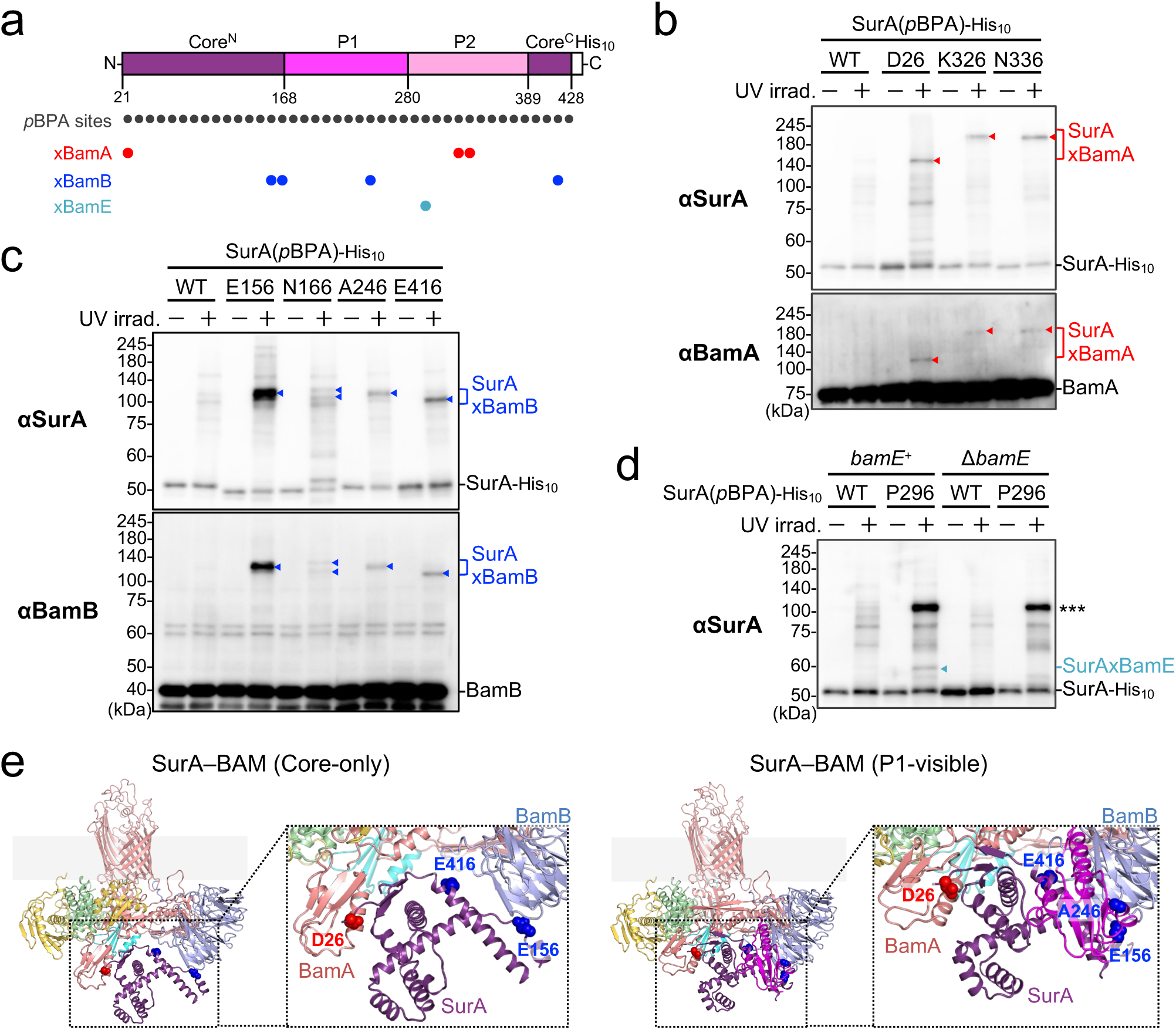
*In vivo* photo-crosslinking analysis of SurA with the BAM complex. **a** Summary of *in vivo* photo-crosslinking sites of SurA. Colored circles represent SurA residues that were crosslinked with Bam factors: BamA (red), BamB (blue), and BamE (cyan). **b–d** *In vivo* photo-crosslinking analysis of SurA. (**b**, **c**) Cells of SN305 (Δ*surA*) carrying both pEVOL-pBpF and pHM1021-*surA(amb)-his10* were grown at 30 °C in LB medium containing 0.5 mM *p*BPA until early log phase and induced with 1 mM IPTG for 1 h to express the indicated SurA(*p*BPA) variants. Cultures were then divided into two portions treated with or without UV irradiation for 10 min at 4 °C. Total cellular proteins were acid-precipitated and analyzed using SDS–PAGE followed by immunoblotting using the indicated antibodies. (**d**) Cells of RM5457 (Δ*surA*) or RM5475 (Δ*surA*, Δ*bamE*) carrying both pEVOL-pBpF and pHM1021-*surA(amb)-his10* were and analyzed in (**b**). *** indicates a nonspecific band that should be a SurA dimer crosslinked product. **e** Mapping of the crosslinking sites on the SurA–BAM cryo-EM structures. SurA residues crosslinked with BamA and BamB are indicated by red and blue spheres, respectively.

Because SurA has been shown to interact with the nonessential Bam factors BamB and BamE, we examined their functional relationship. We constructed Δ*surA* Δ*bamB*, Δ*surA* Δ*bamC*, and Δ*surA* Δ*bamE* double mutant strains. Among these, the Δ*surA* Δ*bamB* and Δ*surA* Δ*bamE* strains exhibited growth defects and higher sensitivity to erythromycin, an antibiotic commonly used to assess OM integrity, compared with the Δ*surA*, Δ*bamB*, or Δ*bamE* single mutants (**Supplementary Fig. 5a**). These results are consistent with previous studies reporting synthetic phenotypes between Δ*surA* and mutations in *bamB* or *bamE*^34,35^. Importantly, BamA levels were unchanged in these strains (**Supplementary Fig. 5b**). These findings indicate that SurA functionally cooperates with BamB and BamE. To further clarify the relationship between SurA and Bam factors, we examined the SurA–BamA photo-crosslinking in the absence of BamB, BamC, or BamE. Deletion of *bamB* or *bamE* significantly reduced formation of the SurA–BamA crosslinked products, despite comparable BamA levels across all strains (**Supplementary Fig. 5c**), with the strongest effect observed in the Δ*bamE* background. Taken together, these results indicate that the interactions with BamB and BamE are crucial for SurA function.

### Cryo-EM structures of P2-visible and P1/P2-visible SurA–BAM complexes

The P2 domain of SurA, which interacts with BamE as shown by our photo-crosslinking analyses (**Fig. 2a, d**), is involved in SurA function, for which BamE has been suggested to play an important role. However, the P2 domain was not observed in either the Core-only or the P1-visible SurA–BAM structures (**Fig. 1a**)^16,17^. Because the molecular mechanism and significance of the interaction between BamE and the P2-domain of SurA remain unclear, we determined the structures of the SurA–BAM complex in which the P2 domain is visible. Based on previous findings^33^, we introduced disulfide crosslinking between the P2 domain and the BAM complex to stabilize their interactions. Cells expressing the combination of SurA(N336C) and BamA(S274C) generated a clear SurA–BamA disulfide crosslinked product (**Supplementary Fig. 6a**). Furthermore, SurA–BamE disulfide crosslinks were formed in all combinations of SurA P296C or I297C with BamE K45C, Q54C, or Y57C (**Supplementary Fig. 6b**). Thus, we introduced either N336C–BamA S274C or SurA I297C–BamE Y57C disulfide crosslinking to stabilize the interaction of the SurA P2 domain with the BAM complex, purified the complexes, and performed cryo-EM single-particle analysis (**Supplementary Fig. 7** and **Supplementary Fig. 8**).

We determined four cryo-EM structures from two disulfide-crosslinked SurA–BAM constructs (**Fig. 3a, b; Supplementary Fig. 9a, b**). These structures can be classified into two conformational states, hereafter referred to as the P2-visible and P1/P2-visible structures. The two structures derived from the different constructs within each state were essentially identical, with Cα RMSD values of 2.33 Å and 2.52 Å for the P2-visible and P1/P2-visible states, respectively (**Supplementary Fig. 10a, b**). In the P2-visible structures, the P2 domain was clearly resolved, whereas the P1 domain was largely unresolved. Based on the PDB entries, the P1/P2-visible structure is the first SurA–BAM structure in which all three domains of SurA are visualized together. In this structure, the P1 domain adopts a compact conformation and is positioned against the SurA Core domain. In both conformational states, the SurA P2 domain contacts BamA and BamE (**Supplementary Fig. 10c–f**). Consistent with our photo-crosslinking analyses (**Fig. 2a**), K326 and N336 residues of SurA are located near BamA, and P296 is positioned close to BamE. Furthermore, we introduced *p*BPA into residues of BamE located at the interface with the SurA P2 domain and performed photo-crosslinking experiments. As expected, BamE–SurA crosslinked products were detected (**Supplementary Fig. 11a**). These photo-crosslinking results strongly support the idea that the SurA P2 domain interacts with BamE as observed in the P2-visible and P1/P2-visible structures. Although the P2-visible and P1/P2-visible structures were stabilized by disulfide crosslinking, our *in vivo* photo-crosslinking analyses demonstrated that similar interactions occur between the SurA P2 domain and BamE under physiological conditions, suggesting that these conformations are not artifacts of crosslinking.

**Fig. 3.**
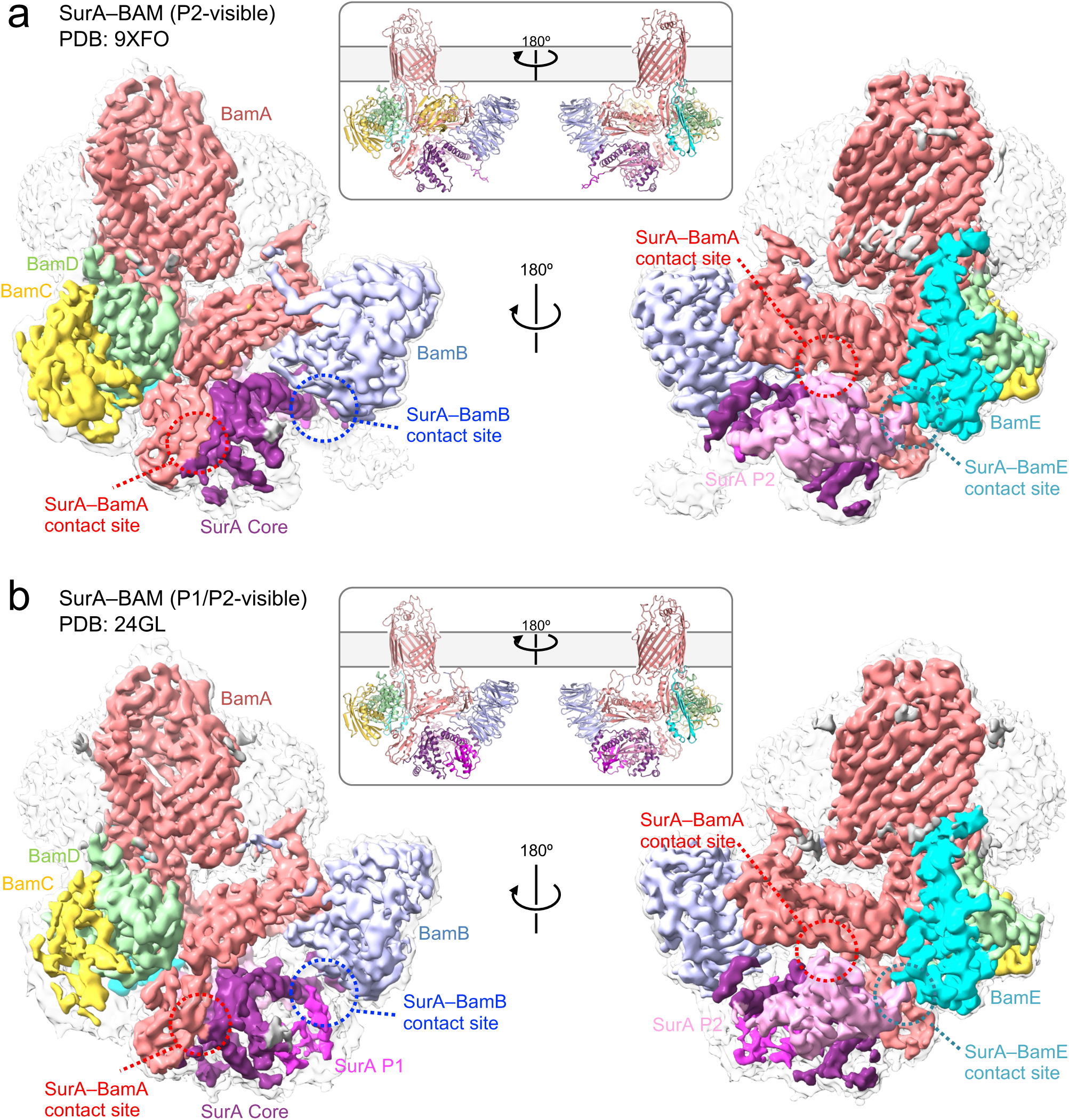
Cryo-EM structures of the P2-visible and P1/P2-visible SurA–BAM complex. **a** Cryo-EM map and cartoon model of the SurA–BAM P2-visible complex (PDB: 9XFO). **b** Cryo-EM map and cartoon model of the SurA–BAM P1/P2-visible complex (PDB: 24GL). The SurA Core, P1 and P2 domains are colored in purple, magenta and light pink, respectively. BamA, BamB, BamC, BamD, and BamE are colored as in Fig. 1. Additional gray map near SurA may be a map from its C-terminal His-tag. Contact sites of SurA with BamA, BamB and BamE are shown in red, blue and cyan circles, respectively.

Next, we superimposed four distinct SurA–BAM structures: the Core-only, P1-visible, P2-visielbe and P1/P2-visible forms (**Fig. 4**). In all four structures, the interaction between the N-terminal region of the SurA Core domain and the POTRA1 domain of BamA is retained. Notably, the position of the SurA Core domain differed among the four structures and was progressively closer to the BAM complex in the order of P1-visible, Core-only, P2-visible, and P1/P2-visible, whereas the other BAM components were well aligned. To examine whether a dynamic conformational change also occurs in living cells, we performed photo-crosslinking analysis using BamB(*p*BPA) derivatives. We introduced *p*BPA into residues of BamB facing the interior of the BAM complex, carried out crosslinking experiments, and identified new SurA-crosslinking sites of BamB (**Supplementary Fig. 11b**). Among these, several sites could contact SurA only in the P2-visible or P1/P2-visible structures, suggesting that the SurA–BAM complex undergoes a conformational change toward a P2-visible- or P1/P2-visible-like state *in vivo*. This conformational change may be mediated through the interaction between the SurA P2 domain and the BAM complex.

**Fig. 4.**
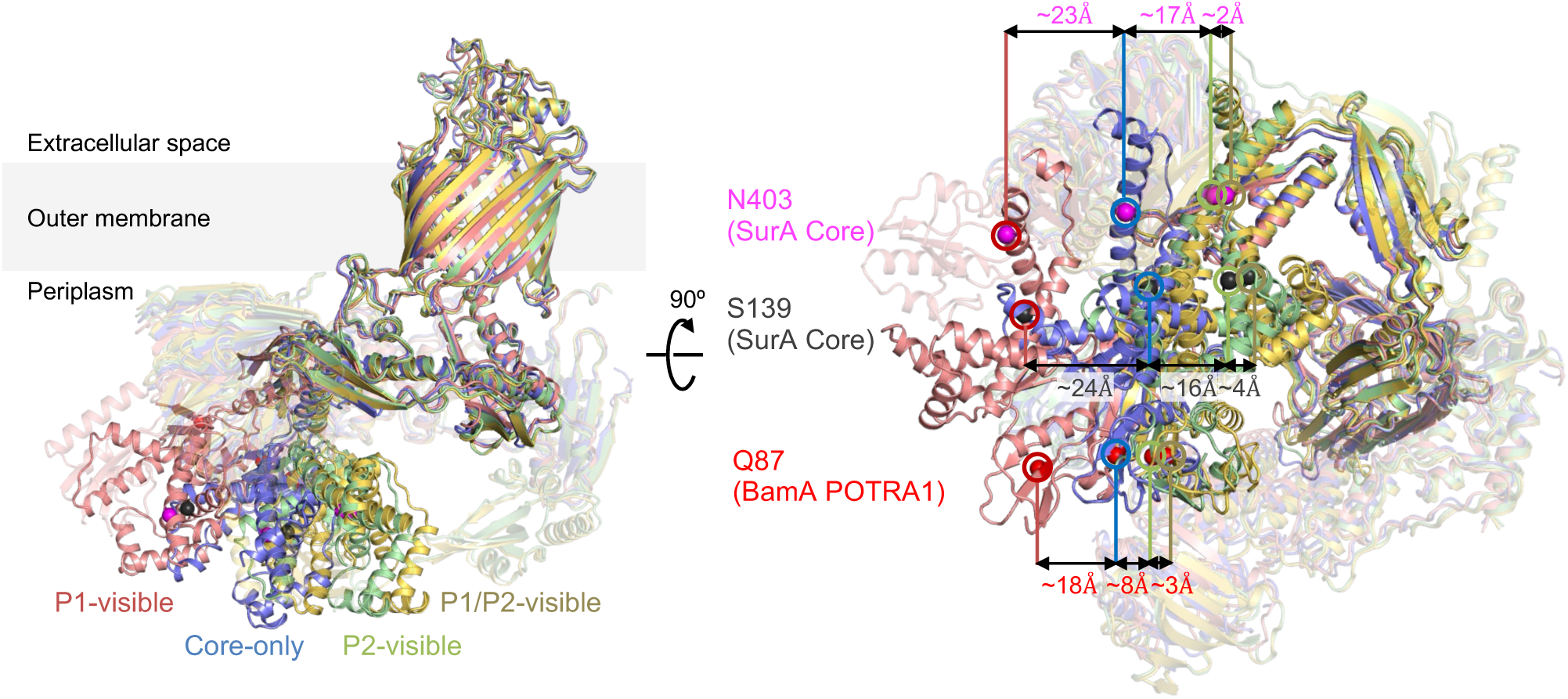
Dynamic relocation of the SurA Core domain among SurA-BAM structures. Cryo-EM structures of the SurA–BAM complex are superposed: P1-visible (red, PDB: 9XBY), Core-only (light blue, PDB: 9XD7), P2-visible (light green, PDB: 9XFO), and P1/P2-visible (yellow, PDB: 24GL). Distances between the S139 and N403 Cα positions in the SurA Core domain and between the Q87 Cα positions in the BamA POTRA1 domain are indicated for each structure.

### The interaction between the SurA P2 domain and BamE is crucial for SurA function

In the P2-visible and P1/P2-visible structures, the P2 domain of SurA interacts with both BamA and BamE. BamA appears to form a transient contact with SurA, whereas BamE is tightly associated with the P2 domain. In fact, residues P296 and I297 in the P2 domain are sandwiched by the Q54 and Y57 residues of BamE, forming a stable interaction (**Fig. 5a**). To examine the functional significance of this SurA–BAM interaction, we constructed two SurA mutants, one lacking residues P296 and I297 (ΔP296–I297) and the other substituting these residues with glutamate (P296E/I297E). We expressed these SurA mutants in the Δ*surA* Δ*bamB* mutant cells and examined their functions (**Fig. 5b**). Although both mutants accumulated to levels comparable to that of wild-type SurA, they exhibited reduced activities, similar to the ΔP2-domain mutant. These results indicate that the BamE-interacting residues are essential for the function of the P2 domain.

**Fig. 5.**
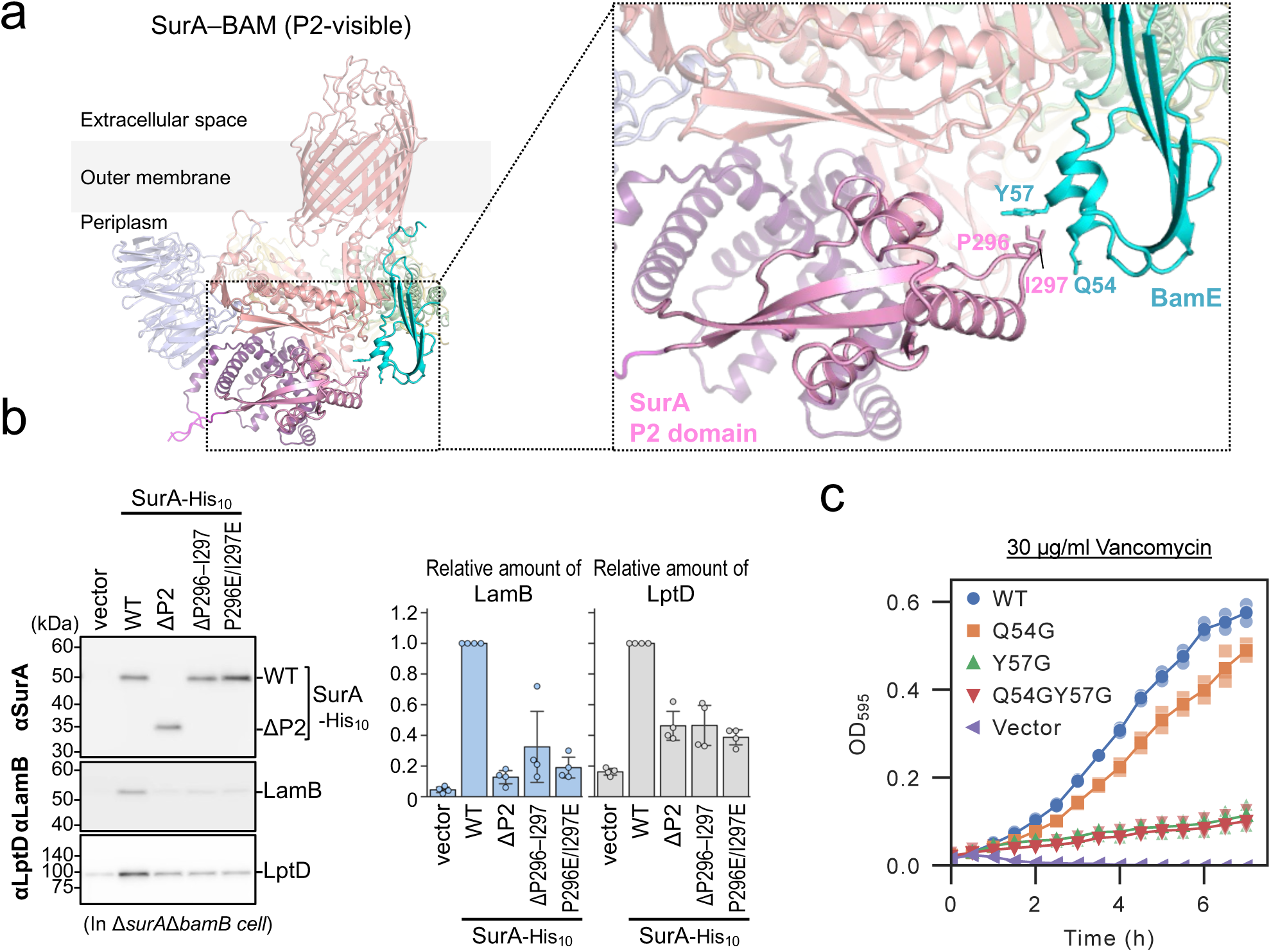
The SurA–BamE interaction is crucial for SurA function. **a** Interface between SurA and BamE. Side chains of SurA and BamE at the contact site are highlighted. **b** Mutational analysis of the BamE-interacting site of SurA. RM5477 (Δ*surA*, Δ*bamB*) cells carrying pHM1021 (vector) or pHM1021-*surA(mut)-his10* plasmids were grown and analyzed as in Fig. 1c. The amounts of LamB and LptD are quantified and shown relative to those of wild-type SurA-His10 expressing cells (n = 4, ± S.D.). **c** Mutational analysis of the SurA-interacting site of BamE. BL21(DE3) Δ*bamE* cells carrying pTnT (vector) or pTnT-*bamE(mut)-his8* were grown in LB medium at 37 °C until log phase, diluted with LB medium supplemented with vancomycin, and cultured. OD595 was monitored at 30-min intervals (n = 3).

In contrast, we introduced glycine substitutions into residues Q54 and Y57 of BamE (**Fig. 5c**). Δ*bamE* cells exhibited higher sensitivity to vancomycin than WT cells due to the disruption of OM integrity. We overexpressed WT BamE or a BamE mutant in Δ*bamE* cells and monitored their growth in medium containing vancomycin. Overexpression of WT BamE complemented the growth defect, whereas Y57G and Q54G Y57G mutants did not. These observations further highlight the functional importance of the interaction between the SurA P2 domain and BamE.

## Discussion

SurA is a major periplasmic chaperone crucial for OMP biogenesis, and is thought to mediate substrate delivery to the BAM complex. Previous biochemical and structural studies elucidated how SurA recognizes unfolded substrates and highlighted the importance of its Core, P1, and P2 domains. More recently, cryo-EM studies of the SurA–BAM complex have provided initial insights into their interaction ^16,17^. However, the molecular mechanism by which SurA delivers substrates to the BAM complex has remained elusive. In this study, we demonstrate that the flexible P1 and P2 domains of SurA play roles as regulators of its Core domain function (**Fig. 1**). Furthermore, our cryo-EM analyses resolved four distinct SurA–BAM structures (Core-only, P1-visible, P2-visible, and P1/P2-visible), revealing multiple conformational states of SurA bound to the BAM complex (**Figs. 1**, **3** and **4**). Crosslinking and functional analyses further indicate that these SurA–BAM architectures are formed *in vivo* (**Fig. 2**) and that the SurA–BAM interactions are important for efficient OMP assembly (**Fig. 5**).

In the BAM complex, BamA and BamD are essential for cell viability, and their functions have been well characterized^8,10–13^. In contrast, the roles of the other Bam components, BamB, BamC, and BamE, are poorly understood. Our crosslinking and structural analyses revealed the detailed interaction modes of SurA with BamB and BamE (**Figs. 1–3**). Moreover, BamB and BamE functionally interact with SurA and contribute to stabilizing the SurA–BAM complex (**Supplementary Fig. 5**). Among them, the interaction between the SurA P2 domain and BamE is particularly important for efficient OMP assembly (**Fig. 5**). This SurA–BamE interaction may promote dynamic conformational changes of SurA on the BAM complex, thereby facilitating substrate delivery. These findings raise the possibility that accessory Bam components not only function within the BAM complex itself but also cooperate with other periplasmic factors, such as SurA, Skp, or BepA, to assist OMP biogenesis. Although BamC has been proposed to undergo conformational changes coupled with those of SurA^17^, we did not observe any physical or functional interactions between them, and its possible involvement in SurA function remains to be carefully evaluated.

The position of the SurA Core domain, which is essential for SurA function, differs markedly among our four SurA–BAM cryo-EM structures (**Fig. 4**). The Core domain is positioned progressively closer to the BAM complex in the order of P1-visible, Core-only, P2-visible, and P1/P2-visible. These findings suggest that SurA bound to the BAM complex may adopt successive conformational states along the substrate delivery pathway. Comparison with the BAM structure lacking SurA further suggests that the BAM conformations observed in our substrate-free SurA–BAM structures are compatible with pre-existing structural states of BAM, given the intrinsic flexibility of the POTRA domains (**Supplementary Fig. 12**).

The SurA Core domain is thought to primarily recognize substrates, with additional contributions from the P1 domain^26,28,29^. Recent cryo-EM studies have also resolved SurA–BAM structures in the presence of substrate proteins^16^. In some of these structures (Arrival complex and Handover complex), substrate is bound to the SurA Core domain (**Supplementary Fig. 13a**). Notably, structures without substrate bound to SurA adopt Core-only-like or P1-visible-like conformations, whereas structures with substrate bound to SurA adopt Core-only-like conformations (**Supplementary Fig. 13**). These observations suggest that substrate binding to the SurA Core domain may shift the SurA-BAM to Core-only-like conformations. When a substrate polypeptide from the previously reported Handover SurA–BAM structure is superposed onto our structures (**Supplementary Fig. 14**), the Core-only and P2-visible structures appear compatible with substrate binding, whereas the P1-visible and P1/P2-visible structures appear less compatible because the P1 domain partially covers the Core domain and would sterically hinder substrate binding. Based on these comparisons, we consider the Core-only and P2-visible structures to be more consistent with substrate-engaged states, whereas the P1-visible and P1/P2-visible structures are more consistent with substrate-free states, such as a substrate-waiting or released state.

Based on this structural information, we propose a working model for SurA-mediated substrate delivery to the BAM complex (**Fig. 6**). After associating with the BAM complex, SurA may adopt substrate-free conformations represented by the P1-visible and Core-only structures. Because its compact P1 arrangement appears to inhibit substrate access, the P1-visible structure is more consistent with a substrate-waiting state. As previously reported^16,27,28^, substrate binding to SurA may trigger a conformational transition of SurA, likely leading to the formation of the Core-only structure representing an early substrate-bound conformation. This transition would bring the Core domain of SurA and its bound substrate closer to the BAM complex. Subsequently, interaction of the P2 domain with the BAM complex, mainly with BamE, may promote or stabilize the P2-visible state, in which the substrate-bound Core domain is positioned closer to the BAM complex. The P1/P2-visible structure may represent a substrate-released state, because the P1 domain adopts a compact arrangement that partially covers the Core domain and appears less compatible with substrate binding in this conformation. However, it remains unclear whether the P1/P2-visible structure represents a state immediately before or after substrate release. This model is reasonable because the substrate-bound SurA Core domain approaches the substrate recognizing regions in the BAM complex, the lateral gate of BamA in its β-barrel domain and the concave surface of the BamD TPR domain^8,10–13^, during substrate transfer to the BAM complex. A substrate delivered by SurA to the BAM complex is finally assembled into the OM via the BAM complex.

**Fig. 6.**
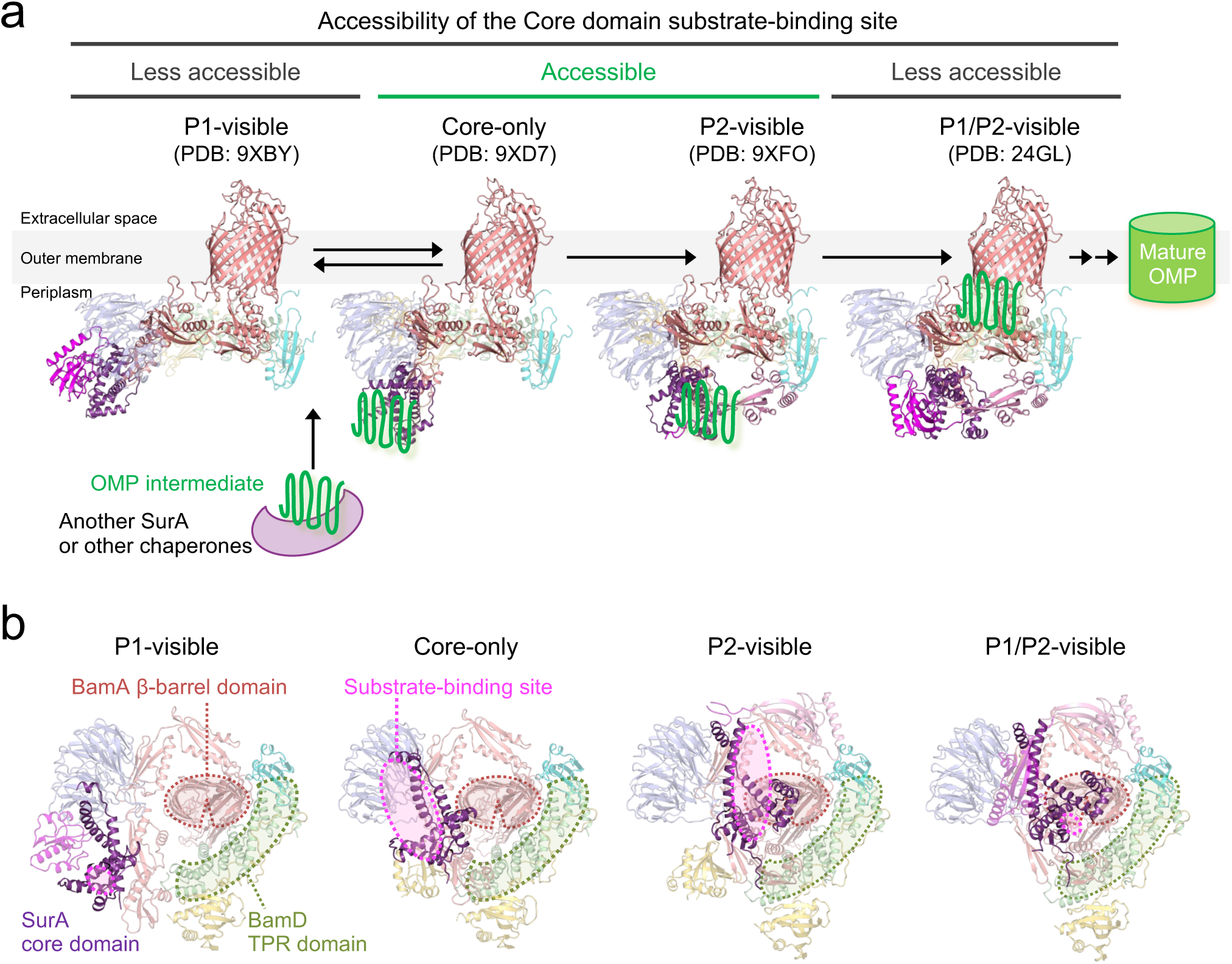
Model for SurA-mediated substrate delivery into the BAM complex. **a** Proposed model of SurA-mediated substrate delivery into the BAM complex. P1-visible, Core-only, P2-visible, and P1/P2-visible conformations may represent successive steps in the substrate delivery pathway, leading to the outer membrane integration and assembly of mature OMPs. **b** Substrate-binding sites in the SurA-BAM complexes viewed from the periplasm. The substrate-binding site of the SurA Core domain is indicated by a pink enclosure. Substrate recognition sites on the BAM complex, BamA and BamD, are indicated by red and light green enclosures, respectively.

Our structural and biochemical analyses have revealed key aspects of the SurA–BAM-mediated substrate delivery mechanism. However, the detailed architecture of SurA–BAM-engaged substrate intermediates and how substrates dynamically interact with SurA and BAM remain to be elucidated. In recently determined substrate-bound SurA–BAM cryo-EM structures, the C-terminal region of the substrate forming a hybrid β-barrel with BamA was clearly resolved, whereas the substrate region associated with SurA was not well defined^16^. This structure may represent a late stage of SurA–BAM-mediated substrate delivery. In this state, SurA may interact with the N-terminal region of the substrate β-barrel domain, similar to the BAM-associated periplasmic protease BepA^36^. Considering that the C-terminal regions of substrate OMPs are recognized by BamA and BamD, it is reasonable to assume that SurA preferentially recognizes the N-terminal regions. SurA has been shown to recognize aromatic residue-enriched sequences within unfolded substrates through its Core and P1 domains^26–28^. Because such sequences are distributed across multiple regions of OMPs, SurA–BAM complexes may contact several parts of a substrate during the early stages of recognition and transfer. Further analyses will be crucial for understanding the initial steps of substrate recognition and delivery.

Moreover, the upstream process by which nascent OMPs are transferred to the SurA–BAM complex is not fully understood. Although SurA has long been thought to directly receive substrates from the Sec translocon and deliver them to BAM, recent structural studies suggest that SurA may associate with BAM and await substrate access^16,17^. One possibility is that other SurA molecules deliver substrates to the SurA–BAM complex, as multiple SurA molecules can bind to a single unfolded substrate^29^. The membrane-anchored periplasmic chaperone complex PpiD/YfgM, which interacts with both the Sec translocon and SurA^32,37^, may contribute to this process. Since PpiD/YfgM assists protein translocation across the IM^37^, the complex may also facilitate substrate transfer from the Sec machinery to the SurA–BAM complex.

The OMP biogenesis pathway, particularly the BAM complex, represents an attractive target for the development of novel antibiotics. Indeed, several small-molecule inhibitors targeting the BAM complex have already been developed^18,19,38^. In this study, we resolved multiple SurA–BAM structures and identified key SurA–BAM interfaces that are crucial for OMP assembly. These structural insights not only advance our fundamental understanding of periplasmic chaperone-assisted OMP biogenesis but also provide valuable information for the design of antibacterial agents. In particular, the SurA–BAM interaction sites identified here may serve as promising targets for the development of novel antibiotics that disrupt OMP assembly in Gram-negative bacteria.

## Methods

### Bacterial strains, plasmids, and primers

The *E. coli* strains, plasmids, and primers used in this study are listed in **Supplementary Tables 1, 2** and **3**, respectively. Details of the mutant strains and plasmids constructed in this study are described in the “Construction of mutant strains” and “Plasmid construction” sections.

### Media and bacterial cultures

The cells were grown in LB medium (Nacalai Tesque), with 50 or 100 µg/mL ampicillin (Amp), 20 µg/mL chloramphenicol (Cm), 25 µg/mL kanamycin (Km), 25 µg/mL tetracycline (Tet), and/or 50 µg/mL spectinomycin (Spc) added as appropriate for growing the plasmid-bearing cells and selecting transformants and transductants. Bacterial growth was monitored using a Mini photo 518R (660 nm; TAITEC Co.).

### Construction of mutant strains

RM5457 (AD16, Δ*surA*) was constructed by removing the *kan* cassette from SN305 (AD16, Δ*surA*::*kan*)^39^ using the pCP20 plasmid^40^. RM5477 (AD16, Δ*surA*, Δ*bamB*::*tet*) was constructed by transducing Δ*bamB*::*tet* from SN147^39^ to RM5457. RM5472 (AD16, Δ*surA*, Δ*bamC*::*kan*) and RM5475 (AD16, Δ*surA*, Δ*bamE*::*kan*) were constructed by transducing Δ*bamC*::*kan* from SN531^39^ and Δ*bamE*::*kan* from SN533^39^ to RM5457, respectively. RM4714 (DY330, *kan araC-ParaBAD-bamA*) was constructed following the same procedure used for the construction of strains with a chromosomal C-terminal His10-tagged gene^41^. First, a *kan araC-ParaBAD-bamA* fragment with a sequence identical to the upstream or downstream region of the *bamA* start codon at the respective ends of the fragment, was polymerase chain reaction (PCR)-amplified from RM3433 (a strain carrying a *kan* cassette at the upstream of an *araC-ParaBAD*) using the ara-bamA-F and ara-bamA-R primers. Next, the chromosomal *bamA* locus of the *E. coli* DY330 strain^42^ was replaced with this fragment using the λ-Red recombination system. RM4674 (HM1742, *purC80*::Tn*10*) was constructed by transducing *purC80*::Tn*10* from RM2098^36^ to HM1742^43^. RM4728 (HM1742, *purC80*::Tn*10*, *kan araC-ParaBAD-bamA*) was constructed by transducing *kan araC-ParaBAD-bamA* from RM4714 to RM4674.

### Plasmid construction

pRM1397 (pHM1021-*surA-his10*) was constructed by PCR amplification of the *surA-his10* fragment from JW0052 (a plasmid encoding *surA*)^44^ using the surA-F and surA-h-R primers, and cloning the fragment into the *NcoI-HindIII* site of pHM1021^45^ following restriction digestion. pHM1021-*surA(amb)-his10* (pRM1400–pRM1440), pHM1021-*surA(Cys)-his10* (pRM1764 and pRM1765), pRM1954 (pHM1021-*surA(ΔP296–I297)-his10*) and pRM1955 (pHM1021-*surA(P296E/I297E)-his10*) plasmids were constructed from pRM1397 by site-directed mutagenesis. To obtain pRM1732–pRM1734 (pHM1021-*surA(Δ172–280, Δ281–389, or Δ172–389)-his10*), pHM1021-*surA-his10* fragments lacking the P1 and/or P2 domains were PCR-amplified from pRM1397 using the S-dP1-F/S-dP1-R, S-dP2-F/S-dP2-R, and S-dP2-F/S-dP12-R primer pairs, respectively. The resulting fragments were self-ligated by *in vitro* recombination using the In-Fusion HD Cloning Kit (Takara Bio).

pRM1399 (pJH113-*surA-ggsg-bamA/bamB/bamC/bamD/bamE-his8*; **Supplementary Fig. 2a**) was constructed as follows: A vector fragment containing *bamA/bamB/bamC/bamD/bamE-his8* was prepared by PCR amplification from pJH114^46^ using the s-bamA-F and s-bamA-R primers. The *surA* fragment was amplified by PCR from JW0052, using the surA-b-F and surA-b-R primers. These two fragments were ligated through *in vitro* recombination using the In-Fusion HD Cloning Kit. pRM1261 (pJH113-*bamA(C690S/C700S)/bamB/bamC/bamD/bamE-his8*) was constructed from pJH114 by two successive rounds of site-directed mutagenesis. The derivatives of pRM1261 carrying an additional Cys mutation, introduced into the *bamA* or *bamE* gene (pRM1713 and pRM1714), were constructed by site-directed mutagenesis. pRM1782 (pJH113-*bamA(S274C/C690S/C700S)/bamB/bamC/bamD/bamE/surA(N336C)-his10*; **Supplementary Fig. 7a**) and pRM1783 (pJH113-*bamA(C690S/C700S)/bamB/bamC/bamD/bamE(Y57C)/surA(I297C)-his10*; **Supplementary Fig. 8a**) were constructed as follows: A vector fragment containing *bamA/bamB/bamC/bamD/bamE* was amplified by PCR from RM1713 or RM1714 using the pBAM-F and pBAM-R primers. A *surA-his10* fragment was amplified from pRM1764 or pRM1765 using the BAM-surA-F and BAM-surA-R primers. The corresponding *bamA–bamE* and *surA-his10* fragments were ligated by *in vitro* recombination using the In-Fusion HD Cloning Kit.

pRM1639 (pHM1021-*bamB-his10*) was constructed by PCR amplification of the *bamB-his10* fragment from pJH114 using the bamB-F and bamB-h-R primers and cloning the fragment into the *NcoI-HindIII* site of pHM1021 following restriction digestion. pHM1021-*bamB(amb)-his10* plasmids (pRM1647–pRM1657) were constructed from pRM1639 by site-directed mutagenesis.

pRM1641 (pHM1021-*bamE-his10*) was constructed by PCR amplification of the *bamE-his10* fragment from pJH114 using the bamE-F and bamE-h-R primers and cloning the fragment into the *NcoI-HindIII* site of pHM1021 following restriction digestion. pHM1021-*bamE(amb)-his10* (pRM1660–pRM1663) and pHM1021-*bamE(Cys)-his10* (pRM1679–pRM1681) plasmids were constructed from pRM1641 by site-directed mutagenesis.

pRM864 (pUC118-*bamA(C690S/C700S)*) was constructed from pRM823 (pUC118-*bamA*)^36^ by two successive rounds of site-directed mutagenesis. Derivatives of pRM864 carrying an additional Cys mutation, introduced into the *bamA* gene (pRM1695 and pRM1696), were constructed by site-directed mutagenesis.

pRM1690 (pSTD689-*surA*) was constructed as follows: A pSTD689-based vector fragment was PCR-amplified from pRM868 (pSTD689-*bepA(E137Q)-his10*) using the pSTD-H-F and pSTD-H-R pairs. A *surA* fragment was prepared by PCR amplification from pRM1397 using the surA-F2 and surA-R2 primers.

These two fragments were ligated by *in vitro* recombination using an In-Fusion HD Cloning Kit to obtain pRM1690. pSTD689-*surA(Cys)* plasmids (pRM1699, pRM1707, pRM1708, and pRM1710) were constructed from pRM1690 by site-directed mutagenesis.

pTnT-*bamE(Gly)-his8* and pTnT-*bamE(Q54G/Y57G)-his8* were constructed from pTnT-bamE-his8^15^ by sequential site-directed mutagenesis.

### Immunoblotting analysis

Acid-denatured proteins, prepared as described in the figure legends for each experiment, were solubilized in SDS-sample buffer (62.5 mM Tris–HCl [pH 6.8], 2% SDS, 10% glycerol, and 0.01% [w/v] bromophenol blue) with or without 10% β-mercaptoethanol (ME). The samples were incubated at 98 °C for 5 min, separated by SDS‒PAGE, and electroblotted onto PVDF membranes (Merck Millipore). The membranes were first blocked with 1% skim milk in phosphate-buffered saline with Tween 20 (PBST) and incubated with anti-SurA^47^ (1/50,000, 1/10,0000, or 1/150,000 dilution), anti-LamB (NBRP; 1/50,000), anti-LptD (1/50,000)^39^, anti-BamA^47^ (1/100,000), anti-BamB^47^ (1/10,000) or anti-His-tag pAb (PM032; MBL; 1/25,000) antibodies. After washing with PBST, the membrane was incubated with a horseradish peroxidase (HRP)-conjugated secondary antibody (goat anti-rabbit IgG (H + L)-HRP conjugate; Bio-Rad; 1/5,000) in PBST. After washing with PBST, the proteins were visualized using Chemi-Lumi One (Nacalai Tesque) or Chemi-Lumi One Super (Nacalai Tesque) and the FUSION Solo S (VILBER) chemiluminescence image analyzer.

### Purification of the SurA–BAM complexes

For purification of the fused SurA–BAM complex, BL21(DE3) (Novagen) cells harboring pRM1399 (**Supplementary Fig. 2**) were inoculated into LB medium supplemented with 0.4% glucose and 50 μg/mL ampicillin and incubated at 37 °C for 16 h. The overnight culture was then inoculated with LB medium containing 50 μg/mL ampicillin and grown at 30 °C until the OD600 reached 0.8–0.9. Isopropyl β-D-thiogalactopyranoside (IPTG) was added at a concentration of 0.5 mM. The cells were further cultured at 37 °C for 1.5 h to induce the SurA–BAM complex expression. Cells were collected by centrifugation, suspended in buffer (20 mM Tris–HCl [pH 8.0], 150 mM NaCl, 1 mM EDTA-Na [pH 8.0], and 0.1 mM PMSF), and disrupted using a Microfluidizer Processor M-110EH at 100 MPa (Microfluidics International). After removing the unbroken cells and protein aggregates by centrifugation, the supernatant was subsequently ultracentrifuged at 40,000 rpm for 60 min at 4 °C (Beckman 45Ti rotor) to isolate the membrane fraction. The membrane fraction was resuspended in solubilization buffer (50 mM Tris–HCl [pH 8.0], 150 mM NaCl, 1% [w/v] n-dodecyl β-maltoside [DDM], 10 mM imidazole-HCl [pH 8.0], and 0.1 mM PMSF) and solubilized by gentle stirring at 4 °C for 60 min. After removing of the insoluble fraction by ultracentrifugation at 40,000 rpm for 30 min at 4 °C (Beckman 45Ti rotor), the supernatant was mixed with 2.5 ml of Ni-NTA agarose resin (QIAGEN), pre-equilibrated with solubilization buffer, and gently stirred at 4 °C for 60 min. Next, 2.5 mL of wash buffer (50 mM Tris–HCl [pH 8.0], 150 mM NaCl, 0.05% [w/v] DDM, 50 mM imidazole-HCl [pH 8.0], and 0.1 mM PMSF) was added to the column three times. Then, 2.5 mL of elution buffer (50 mM Tris–HCl [pH 8.0], 150 mM NaCl, 0.05% [w/v] DDM, 500 mM imidazole-HCl [pH 8.0], and 0.1 mM PMSF) was added to the column six times. The eluted fractions containing SurA-BamA, BamB, BamC, BamD, and BamE-His8 were collected and concentrated using an Amicon Ultra 50K NMWL (Merck Millipore). After ultracentrifugation at 45,000 rpm for 30 min at 4 °C (Himac S55A2 rotor), the concentrated sample was loaded onto a Superose^TM^ 6 Increase 10/300 GL column (Cytiva) equilibrated with SEC buffer (50 mM Tris–HCl [pH 8.0], 150 mM NaCl, 0.02% [w/v] DDM, and 0.1 mM PMSF). The fractions eluted with the target proteins were collected and further concentrated using an Amicon Ultra 50K NMWL.

For purification of the disulfide-crosslinked SurA–BAM complexes. *E. coli* BL21(DE3) cells carrying pRM1782 (**Supplementary Fig. 7**) or pRM1783 (**Supplementary Fig. 8**) were cultured and harvested as described above. The collected membrane fractions were solubilized and purified by Ni-NTA affinity chromatography. The eluted samples were concentrated and further purified by size-exclusion chromatography using a Superdex^TM^ 200 Increase 10/300 GL column (Cytiva) equilibrated with SEC buffer. Fractions containing the disulfide-crosslinked SurA–BAM complexes were collected and concentrated.

### Cryo-EM grid preparation

Quantifoil holey carbon grids (Cu R1.2/R1.3, 300 mesh) were glow-discharged at 7 Pa and 10 mA for 10 seconds using a JEC-3000FC sputter coater (JEOL) before sample application. For the disulfide-crosslinked SurA–BAM complex, Quantifoil holey carbon grids (Cu R1.2/R1.3, 300 mesh) were glow-discharged for 45 s at 20 mA using a GloQube (Quorum) instrument. A 3-μL aliquot of the SurA–BAM complexes (∼15 mg/mL) was applied onto the grids, which were blotted for 2 or 3 seconds at 100% humidity and 8 °C with a blot force of 10 or 4, and then plunged into liquid ethane using a Vitrobot Mark IV (Thermo Fisher Scientific).

### Cryo-EM data collection and processing

The cryo-EM datasets were collected using a CRYO ARM 300 transmission electron microscope (JEOL) operated at an accelerating voltage of 300 kV, equipped with a cold-field emission gun, an in-column Omega-type energy filter, and a Gatan K3 camera (Gatan) at SPring-8. Images were collected at a nominal magnification of 60,000×, corresponding to a calibrated pixel size of 0.752 Å/pixel, 50 frames per image with a total exposure dose of 50 e−/Å2 using a SerialEM^48^. For the disulfide-crosslinked SurA–BAM complex, the cryo-EM dataset was collected on a Krios G4 Cryo-TEM instrument operated at an accelerating voltage of 300 kV and equipped using a Selectris X energy filter and Falcon4i direct electron detector (Thermo Fisher Scientific) at National Research and Innovation (BRIN) in Indonesia. Images were collected at a nominal magnification of 165,000×, corresponding to a calibrated pixel size of 0.76 Å/pixel, 40 frames per image with a total exposure dose of 30 e−/Å2, using the EPU software (Thermo Fisher Scientific).

Data processing was performed using the CryoSPARC v4.5.3–v4.6.2 software platform^49^. Cryo-EM data-processing for various SurA–BAM complexes was performed by the procedures shown in **Supplementary Figs. 2d, 7d,** and **8d**. The acquired movies were aligned using patch motion correction, and the contrast transfer function (CTF) parameters were estimated using the Patch CTF estimation. The particles were initially auto-picked from 200 micrographs using the Blob Picker, and the resulting particles were subjected to 2D classification. The obtained 2D class averages were then used as templates for automatic particle picking with the Template Picker. Particles were extracted with a downsampling via Fourier cropping and subjected to 2D classification. Next, these selected particles were subjected to ab initio 3D reconstruction to generate initial 3D models. The particles were then subjected to several rounds of heterogeneous refinement to remove junk particles using the initial models as references. The best 3D class particles were re-extracted at the original pixel size. These particles were subjected to global/local CTF refinement and Non-Uniform (NU) refinement, which resulted in final cryo-EM maps.

For the dataset of the fused SurA–BAM complex, to analyze structural heterogeneity, the refined 295,437 particles were subjected to two rounds of 3D classification without alignment in CryoSPARC. Each particle subset was subjected to NU refinement, resulting in two final maps: a Core-only SurA–BAM complex and a P1-visible SurA–BAM complex (**Supplementary Fig. 2d**). For the datasets of the disulfide-crosslinked SurA–BAM complexes 1 and 2, the refined 742,649 and 657,643 particles, respectively, were also subjected to one or two rounds of 3D classification without alignment in CryoSPARC. Each particle subset was then subjected to NU refinement, yielding two final maps for each dataset: a P2-visible SurA–BAM complex and a P1/P2-visible SurA–BAM complex (**Supplementary Figs. 7d** and **8d**).

### Model building and refinement

The SurA–BAM structures (PDB ID: 8PZ2 and 8PZ1) and the the AlphaFold2/3-predicted model^50,51^ were docked into the obtained cryo-EM maps using UCSF Chimera X^52^. The fitted structures were manually adjusted in Coot^53^, followed by further refinements using the phenix.real_space_refine in PHENIX^54^. The data processing and refinement statistics are summarized in **Supplementary Table 4**. The molecular model and cryo-EM map were visualized using PyMOL (https://pymol.org/) or UCSF ChimeraX.

### *In vivo* photo-crosslinking analysis

pEVOL-pBpF expresses an evolved tRNA/aminoacyl-tRNA synthetase pair that enables the *in vivo* incorporation of *p*BPA into an amber codon site of a target protein via *amber* suppression^55,56^. UV irradiation to a cell expressing a *p*BPA-incorporated protein induces the formation of covalent crosslinking products between *p*BPA in the target protein and a nearby protein, which allows for the detection of their *in vivo* interaction^55–57^. Cells were grown at 30 °C in LB medium supplemented with 0.5 mM *p*BPA and 0.02% arabinose until the early log phase, followed by induction with 1 mM IPTG for 1 h. Half of the cell cultures was transferred to a Petri dish and UV-irradiated at 4 °C for 10 min using a B-100AP UV lamp (365 nm; UVP, LLC.) at a distance of 4 cm. The other half was placed on ice as a non-UV-irradiated samples. Total cellular proteins were precipitated with 5% trichloroacetic acid (TCA), washed with acetone, solubilized in SDS sample buffer containing ME, and incubated at 98 °C for 5 min. Samples were then analyzed by SDS‒PAGE and immunoblotting.

### Disulfide crosslinking

For SurAxBamA disulfide crosslinking, RM4728 (HM1742, *purC80*::Tn*10*, *kan araC-ParaBAD-bamA*) cells carrying a combination of plasmids encoding WT or a Cys-introduced mutant of BamA and SurA were inoculated into LB medium containing 0.1% arabinose at 30 °C for 16 h. The cultured cells were washed and resuspended in LB medium. The cell suspensions were inoculated into LB medium supplemented with 1 mM IPTG at 30 °C for 3 h. Total cellular proteins were precipitated with 5% TCA, washed with acetone, and suspended in SDS sample buffer containing 12.5 mM NEM to block the free SH groups of the Cys residues. Half of the samples were mixed with 10% ME, incubated at 98 °C for 5 min. Samples were analyzed by 7.5% Laemmli SDS‒PAGE and immunoblotting using anti-SurA and anti-BamA antibodies. For SurAxBamE disulfide crosslinking, RM5475 (AD16, Δ*surA*, Δ*bamE*::*kan*) cells carrying a combination of plasmids encoding WT or a Cys-introduced mutant of SurA and BamE-His10 were inoculated into LB medium containing 0.4% glucose at 30 °C for 16 h. The cultured cells were inoculated into LB medium supplemented with 1 mM IPTG at 30 °C for 2.5 h. Total cellular proteins were precipitated with 5% TCA, washed with acetone, and suspended in SDS sample buffer containing 12.5 mM NEM. Half of the samples were mixed with 10% ME, incubated at 98 °C for 5 min. Samples were analyzed by 7.5% Laemmli SDS‒PAGE or 10% wide-range gel and immunoblotting using anti-SurA and anti-His antibodies.

## Acknowledgements

We thank Kayo Abe for secretarial assistance, Kunihito Yoshikaie for technical support, Ian Smith for English editing, Harris D. Bernstein for kindly providing pJH114, and the scientists of SPring-8 Structural Biology Beamlines for helping with data collection. ChatGPT (OpenAI) was used to improve the English of this manuscript. The cryo-EM experiments were performed at SPring-8 in Japan with the approval of the Japan Synchrotron Radiation Research Institute (proposal nos. 2024A2742 and 2024A2759) and at National Research and Innovation Agency (BRIN) in Indonesia. Optimization of sample and grid preparation conditions was performed using a Glacios cryo-transmission electron microscope with the technical support of the Cryo-EM Facility, Institute for Life and Medical Sciences, Kyoto University in Japan. This work was supported by JSPS/MEXT KAKENHI (Grant Nos. JP21KK0126, JP22K15061, JP22H05567, JP24KK0138, JP25K00267 to R.M., Grant Nos. JP23K14146, JP25K18422 to H.K., Grant Nos. JP24KK0138 to T.S., and Grant Nos. JP25K02226, JP22H02567, JP22H02586, JP21H05155, JP21H05153, JP21K19226, JP21KK0125 to T.T.), the JST FOREST Program (JPMJFR2064 to T.S.), JST NEXUS (Nos. JPMJNX25E5), private research foundations (the Institute for Fermentation (Y-2024-02-006 to R.M., Y-2025-2-046 to H.K., and G-2024-2-034 to T.T., the Chemo-Sero-Therapeutic Research Institute, Takeda Science Foundation to T.T.), JST SPRING (Grant No. JPMJSP2140 to Y.S.T.), and RIIM-KI JST (Grant No. 76/II.7/HK/2025). This research was partially supported by Platform Project for Supporting Drug Discovery and Life Science Research (Basis for Supporting Innovative Drug Discovery and Life Science Research (BINDS)) from AMED under grant number JP25ama121001.

## Author Contributions

Conceptualization: R.M.

Methodology: R.M., H.K., T.S., and T.T.

Investigation: R.M., H.K., N.M., Y.M., W.Y., Y.S.T., D.H.Y.Y., Y.N., H.S., T.S., and T.T.

Visualization: R.M., H.K., Y.M., and T.T.

Supervision: R.M., and T.T.

Writing—original draft: R.M. and T.T.

Writing—review and editing: R.M. and T.T.

## Competing interests

The authors declare no competing interests.

## Data Availability

The coordinates and corresponding electron potential maps have been deposited in the Protein Data Bank (PDB) and Electron Microscopy Data Bank (EMDB) under the following accession codes: Core-only SurA-BAM complex, PDB 9XD7 and EMDB-66757; P1-visible SurA-BAM complex, PDB 9XBY and EMDB-66714; P2-visible SurA-BAM complex, PDB 9XFO and EMDB-66834; P1/P2-visible SurA-BAM complex, PDB 24GL and EMDB-69488; P2-visible SurA-BAM complex 2, PDB 9XFG and EMDB-66821; and P1/P2-visible SurA-BAM complex 2, PDB 24GT and EMDB-69496, respectively. Source data, including uncropped scans of all blots and gels, are provided with this paper.

## Code Availability

This paper does not report original code.

## Supplementary materials

Supplementary Figs. 1 to 14

Supplementary Tables 1 to 4

**Supplementary Fig. 1 |** OMP assembly pathway and previous structures of the SurA–BAM complexes.

**Supplementary Fig. 2 |** Cryo-EM analysis of the fused SurA–BAM.

**Supplementary Fig. 3 |** Structural comparisons of the SurA–BAM complexes.

**Supplementary Fig. 4 |** *In vivo* photo-crosslinking analysis of SurA.

**Supplementary Fig. 5 |** Functional relationship between SurA and Bam factors.

**Supplementary Fig. 6 |** *In vivo* disulfide crosslinking between SurA and the BAM complex.

**Supplementary Fig. 7 |** Cryo-EM analysis of the disulfide crosslinked SurA–BAM complex 1.

**Supplementary Fig. 8 |** Cryo-EM analysis of the disulfide crosslinked SurA–BAM complex 2.

**Supplementary Fig. 9 |** Cryo-EM structures of the P2-visible 2 and P1/P2-visible 2 SurA–BAM complexes.

**Supplementary Fig. 10 |** Structural details of cryo-EM structures of the P2-visible and P1/P2-visible SurA/BAM complexes.

**Supplementary Fig. 11 |** *In vivo* photo-crosslinking between SurA and BamE or BamB.

**Supplementary Fig. 12 |** Comparison of BamA in our SurA–BAM cryo-EM structures with a SurA-lacking BAM structure.

**Supplementary Fig. 13 |** Cryo-EM structures of the SurA–BAM complex.

**Supplementary Fig. 14 |** Substrate binding to the SurA Core domain in SurA-BAM cryo-EM structures.

**Supplementary Table 1:** Strains used in this study.

**Supplementary Table 2:** Plasmids used in this study.

**Supplementary Table 3:** Primers used in this study.

**Supplementary Table 4:** Cryo-EM data collection and refinement statistics.

## Notes

### Competing Interest Statement

The authors have declared no competing interest.

### Summary of Updates

The manuscript has been revised following peer review. Figures, supplementary materials, and text have been updated to address reviewer comments.

