## Supplemental data for "Cryo-EM structures of the BAM–P1/P2-visible SurA complex reveal dynamic and cooperative interactions in outer membrane protein assembly"

**Supplementary Table 1 | Strains used in this study.**

| Strain | Genotype | Reference |
|---|---|---|
| AD16 | Δ*pro-lac thi*/F' $lacI^q$ *ZΔM15* $Y^+$ $pro^+$ | Kihara *et al.* [55] |
| SN305 | AD16, Δ*surA*::*kan* | Narita *et al.* [39] |
| SN147 | AD16, Δ*bamB*::*tet* | Narita *et al.* [39] |
| SN531 | AD16, Δ*bamC*::*kan* | Narita *et al.* [39] |
| SN533 | AD16, Δ*bamE*::*kan* | Narita *et al.* [39] |
| RM5457 | AD16, Δ*surA*::*FRT* | This study |
| RM5477 | AD16, Δ*surA*::*FRT* Δ*bamB*::*tet* | This study |
| RM5472 | AD16, Δ*surA*::*FRT* Δ*bamC*::*kan* | This study |
| RM5475 | AD16, Δ*surA*::*FRT* Δ*bamE*::*kan* | This study |
| MC4100 | $F^-$ *araD139 Δ(argF-lac)U169 rpsL150 relA1 flbB5301 deoC1 ptsF25 rbsR* | Silhavy *et al.* [59] |
| CU141 | MC4100/F' $lacI^q$ $lacZ^+$, $Y^+$, $A^+$ | Akiyama *et al.* [60] |
| HM1742 | CU141, $ara^+$ | Mori and Ito [43] |
| RM4674 | HM1742, *purC80*::Tn*10* | This study |
| RM4728 | HM1742, *purC80*::Tn*10 kan araC-*$P_{araBAD}$*-bamA* | This study |
| DY330 | W3110, Δ*lacU169 gal490 λcI857* Δ(*cro-bioA*) | Yu e*t al.* [42] |
| RM4714 | DY330, *kan araC-*$P_{araBAD}$*-bamA* | This study |
| BL21(DE3) | $F^-$, *ompT, hsdS*$_B$($r_B^-$ $m_B^-$), *gal*(*λc*I 857, *ind*1, *Sam*7, *nin*5, *lacUV*5-T7*gene*1), *dcm*(DE3) | Novagen |
| Δ*bamE* | BL21(DE3)*, Δ*bamE*::*kan* | Thewasano *et al.* [14] |

**Supplementary Table 2 | Plasmids used in this study.**

| Plasmid | Vector | Encoded gene and description | Reference or source |
|---|---|---|---|
| pEVOL-pBpF | | p15A-derivative encoding an evolved *M. jannaschii* aminoacyl-tRNA synthetase/suppressor tRNA pair for incorporation of *p*BPA; $Cm^R$ | Young *et al.* [56] |
| pJH113 | | Expression vector; $P_{trc}$, $Amp^R$ | Roman-Hernandez *et al.* [46] |
| pJH114 | pJH113 | *bamA/bamB/bamC/bamD/bamE-his$_8$* | Roman-Hernandez *et al.* [46] |
| pRM1399 | pJH113 | *surA-ggsg-bamA/bamB/bamC/bamD/bamE-his$_8$* | This study |
| pRM1261 | pJH113 | *bamA(C690S, C700S)/bamB/bamC/bamD/bamE-his$_8$* | This study |
| pRM1713 | pJH113 | *bamA(S274C, C690S, C700S)/bamB/bamC/bamD/bamE-his$_8$* | This study |
| pRM1714 | pJH113 | *bamA(C690S, C700S)/bamB/bamC/bamD/bamE(Y57C)-his$_8$* | This study |
| pRM1782 | pJH113 | *bamA(S274C, C690S, C700S)/bamB/bamC/bamD/bamE/surA(N336C)-his$_{10}$* | This study |
| pRM1783 | pJH113 | *bamA(C690S, C700S)/bamB/bamC/bamD/bamE(Y57C)/surA(I297C)-his$_{10}$* | This study |
| pHM1021 | | Expression vector; $P_{lac}$, $Amp^R$ | Ishii *et al.* [45] |
| pRM1397 | pHM1021 | *surA-his$_{10}$* | This study |
| pRM1400 | pHM1021 | *surA(D26amb)-his$_{10}$* | This study |
| pRM1401 | pHM1021 | *surA(V36amb)-his$_{10}$* | This study |
| pRM1402 | pHM1021 | *surA(M46amb)-his$_{10}$* | This study |
| pRM1403 | pHM1021 | *surA(A56amb)-his$_{10}$* | This study |
| pRM1404 | pHM1021 | *surA(L66amb)-his$_{10}$* | This study |
| pRM1405 | pHM1021 | *surA(M76amb)-his$_{10}$* | This study |
| pRM1406 | pHM1021 | *surA(K86amb)-his$_{10}$* | This study |
| pRM1407 | pHM1021 | *surA(L96amb)-his$_{10}$* | This study |
| pRM1408 | pHM1021 | *surA(Q106amb)-his$_{10}$* | This study |
| pRM1409 | pHM1021 | *surA(S116amb)-his$_{10}$* | This study |
| pRM1410 | pHM1021 | *surA(N126amb)-his$_{10}$* | This study |
| pRM1411 | pHM1021 | *surA(M136amb)-his$_{10}$* | This study |
| pRM1412 | pHM1021 | *surA(V146amb)-his$_{10}$* | This study |
| pRM1413 | pHM1021 | *surA(E156amb)-his$_{10}$* | This study |
| pRM1414 | pHM1021 | *surA(N166amb)-his$_{10}$* | This study |
| pRM1415 | pHM1021 | *surA(L176amb)-his$_{10}$* | This study |
| pRM1416 | pHM1021 | *surA(N186amb)-his$_{10}$* | This study |
| pRM1417 | pHM1021 | *surA(E196amb)-his$_{10}$* | This study |

| | | | |
|---|---|---|---|
| pRM1418 | pHM1021 | *surA(A206amb)-his$_{10}$* | This study |
| pRM1419 | pHM1021 | *surA(A216amb)-his$_{10}$* | This study |
| pRM1420 | pHM1021 | *surA(L226amb)-his$_{10}$* | This study |
| pRM1421 | pHM1021 | *surA(I236amb)-his$_{10}$* | This study |
| pRM1422 | pHM1021 | *surA(A246amb)-his$_{10}$* | This study |
| pRM1423 | pHM1021 | *surA(V256amb)-his$_{10}$* | This study |
| pRM1424 | pHM1021 | *surA(H266amb)-his$_{10}$* | This study |
| pRM1425 | pHM1021 | *surA(E276amb)-his$_{10}$* | This study |
| pRM1426 | pHM1021 | *surA(H286amb)-his$_{10}$* | This study |
| pRM1427 | pHM1021 | *surA(P296amb)-his$_{10}$* | This study |
| pRM1428 | pHM1021 | *surA(K306amb)-his$_{10}$* | This study |
| pRM1429 | pHM1021 | *surA(S316amb)-his$_{10}$* | This study |
| pRM1430 | pHM1021 | *surA(K326amb)-his$_{10}$* | This study |
| pRM1431 | pHM1021 | *surA(N336amb)-his$_{10}$* | This study |
| pRM1432 | pHM1021 | *surA(P346amb)-his$_{10}$* | This study |
| pRM1433 | pHM1021 | *surA(A356amb)-his$_{10}$* | This study |
| pRM1434 | pHM1021 | *surA(S366amb)-his$_{10}$* | This study |
| pRM1435 | pHM1021 | *surA(H376amb)-his$_{10}$* | This study |
| pRM1436 | pHM1021 | *surA(V386amb)-his$_{10}$* | This study |
| pRM1437 | pHM1021 | *surA(R396amb)-his$_{10}$* | This study |
| pRM1438 | pHM1021 | *surA(F406amb)-his$_{10}$* | This study |
| pRM1439 | pHM1021 | *surA(E416amb)-his$_{10}$* | This study |
| pRM1440 | pHM1021 | *surA(L426amb)-his$_{10}$* | This study |
| pRM1764 | pHM1021 | *surA(I297C)-his$_{10}$* | This study |
| pRM1765 | pHM1021 | *surA(N336C)-his$_{10}$* | This study |
| pRM1732 | pHM1021 | *surA(Δ172–280)-his$_{10}$* | This study |
| pRM1734 | pHM1021 | *surA(Δ281–389)-his$_{10}$* | This study |
| pRM1735 | pHM1021 | *surA(Δ172–389)-his$_{10}$* | This study |
| pRM1954 | pHM1021 | *surA(ΔP296–I297)-his$_{10}$* | This study |
| pRM1955 | pHM1021 | *surA(P296E/I297E)-his$_{10}$* | This study |
| pRM1639 | pHM1021 | *bamB-his$_{10}$* | This study |
| pRM1647 | pHM1021 | *bamB(Q170amb)-his$_{10}$* | This study |
| pRM1648 | pHM1021 | *bamB(N186amb)-his$_{10}$* | This study |
| pRM1649 | pHM1021 | *bamB(D188amb)-his$_{10}$* | This study |
| pRM1650 | pHM1021 | *bamB(R215amb)-his$_{10}$* | This study |
| pRM1651 | pHM1021 | *bamB(Q225amb)-his$_{10}$* | This study |

| | | | |
|---|---|---|---|
| pRM1652 | pHM1021 | *bamB(M226amb)-his$_{10}$* | This study |
| pRM1653 | pHM1021 | *bamB(R231amb)-his$_{10}$* | This study |
| pRM1654 | pHM1021 | *bamB(Q234amb)-his$_{10}$* | This study |
| pRM1655 | pHM1021 | *bamB(T236amb)-his$_{10}$* | This study |
| pRM1656 | pHM1021 | *bamB(S238amb)-his$_{10}$* | This study |
| pRM1657 | pHM1021 | *bamB(R243amb)-his$_{10}$* | This study |
| pRM1641 | pHM1021 | *bamE-his$_{10}$* | This study |
| pRM1660 | pHM1021 | *bamE(K45amb)-his$_{10}$* | This study |
| pRM1661 | pHM1021 | *bamE(M50amb)-his$_{10}$* | This study |
| pRM1662 | pHM1021 | *bamE(Q54amb)-his$_{10}$* | This study |
| pRM1663 | pHM1021 | *bamE(Y57amb)-his$_{10}$* | This study |
| pRM1679 | pHM1021 | *bamE(K45C)-his$_{10}$* | This study |
| pRM1680 | pHM1021 | *bamE(Q54C)-his$_{10}$* | This study |
| pRM1681 | pHM1021 | *bamE(Y57C)-his$_{10}$* | This study |
| pUC118 | | Expression vector; $P_{lac}$, $Amp^R$ | Takara Bio |
| pRM823 | pUC118 | *bamA* | Miyazaki *et al.* [36] |
| pRM864 | pUC118 | *bamA(C690S/C700S)* | This study |
| pRM1695 | pUC118 | *bamA(L78C, C690S, C700S)* | This study |
| pRM1696 | pUC118 | *bamA(S274C, C690S, C700S)* | This study |
| pSTD689 | | Expression vector; $P_{lac}$, $Spc^R$ | Kanehara *et al.* [61] |
| pRM1690 | pSTD689 | *surA* | This study |
| pRM1699 | pSTD689 | *surA(V24C)* | This study |
| pRM1707 | pSTD689 | *surA(P296C)* | This study |
| pRM1708 | pSTD689 | *surA(I297C)* | This study |
| pRM1710 | pSTD689 | *surA(N336C)* | This study |
| pTnT | pTnT | Expression vector, $P_{T7\text{-}SP6}$, $Amp^R$ | Promega |
| pTnT-bamE | pTnT | *bamE* | Maruno *et al.* [15] |
| pTnT-bamE-his8 | pTnT | *bamE-his$_8$* | Maruno *et al.* [15] |
| pTnT-bamE-his8 (Q54G) | pTnT | *bamE(Q54G)-his$_8$* | This study |
| pTnT-bamE-his8 (Y57G) | pTnT | *bamE(Y57G)-his$_8$* | This study |
| pTnT-bamE-his8 (Q54G/Y57G) | pTnT | *bamE(Q54G/Y57G)-his$_8$* | This study |

**Supplementary Table 3 | Primers used in this study.**

| Name | Sequence (5' to 3') |
|---|---|
| ara-bamA-F | TTTCTCTCGGTTATGAGAGTTAGTTAGGAAGAACGCATAATAACGCATATGAATATCCTCCTTAG |
| ara-bamA-R | GCTGCTAAACAGCAGCGACGCTATGAGCAACTTTTTCATCGCCATGGTGAATTCCTCCTGCTAG |
| surA-F | GCGCCCATGGCGAAGAACTGGAAAACGCTGC |
| surA-h-R | GCGCAAGCTTAATGATGATGATGATGATGATGATGATGATGGTTGCTCAGGATTTTAACG |
| S-dP1-F | AACTTCGGTCACCGAGCTGGCGTCGTTTTGGTTAC |
| S-dP1-R | TCGGTGACCGAAGTTCATGC |
| S-dP2-F | TTTCTGCGCAGCGTCGATATTTTTGCTTTCGCCGC |
| S-dP2-R | GACGCTGCGCAGAAAGATCG |
| S-dP12-F | TTTCTGCGCAGCGTCGCTGGCGTCGTTTTGGTTAC |
| s-bamA-F | AACGGTGGCTCTGGTGCTGAAGGGTTCGTAGTG |
| s-bamA-R | CATCGTTATTATGCGTTCTTC |
| surA-b-F | CGCATAATAACGATGAAGAACTGGAAAACGCTG |
| surA-b-R | ACCAGAGCCACCGTTGCTCAGGATTTTAACG |
| pBAM-F | GATCCTCTAGAGTCGACCTGCAGGCATG |
| pBAM-R | TTAGTTACCACTCAGCGCAGG |
| BAM-surA-F | CTGAGTGGTAACTAACACACAGGAAACAGACCATG |
| BAM-surA-R | CGACTCTAGAGGATCGGCCAGTGCCAAGCTTAATG |
| bamB-F | GCGCCCATGGCGCAATTGCGTAAATTACTGC |
| bamB-h-R | GCGCAAGCTTAATGATGATGATGATGATGATGATGATGATGACGTGTAATAGAGTACACG |
| bamE-F | GCGCCCATGGCGCGCTGTAAAACGCTGACTG |
| bamE-h-R | GCGCAAGCTTAATGATGATGATGATGATGATGATGATGATGGTTACCACTCAGCGCAGG |
| pSTD-H-F | CTCGAGCATCATCATCATC |
| pSTD-H-R | CATAACCTCTATCCTGTATTTC |
| surA-F2 | AGGATAGAGGTTATGAAGAACTGGAAAACGCTGC |
| surA-R2 | ATCCCCGGGTACTTAGTTGCTCAGGATTTTAACG |

### Supplementary Table 4 | Cryo-EM data collection and refinement statistics.

| dataset | | | | | | |
|---|---|---|---|---|---|---|
| Structure | ***P1-visible*** | ***Core-only*** | ***P2-visible 1*** | ***P1/P2-visible 1*** | ***P2-visible 2*** | ***P1/P2-visible 2*** |
| EMD ID | **(EMD-66714)** | **(EMD-66757)** | **(EMD-66834)** | **(EMD-69488)** | **(EMD-66821)** | **(EMD-69496)** |
| PDB ID | **(9XBY)** | **(9XD7)** | **(9XFO)** | **(24GL)** | **(9XFG)** | **(24GT)** |
| **Data collection and processing** | | | | | | |
| Microscope | CRYO ARM 300 | CRYO ARM 300 | Titan Krios | Titan Krios | CRYO ARM 300 | CRYO ARM 300 |
| Magnification | x60,000 | x60,000 | x165,000 | x165,000 | x60,000 | x60,000 |
| Voltage (kV) | 300 | 300 | 300 | 300 | 300 | 300 |
| Electron exposure (e–/Å$^2$) | 49.92 | 49.92 | 30 | 30 | 50 | 50 |
| Defocus range (μm) | -1.4 to -1.6 | -1.4 to -1.6 | -0.6 to -1.6 | -0.6 to -1.6 | -1.4 to -1.6 | -1.4 to -1.6 |
| Pixel size (Å) | 0.752 | 0.752 | 0.76 | 0.76 | 0.752 | 0.752 |
| Symmetry imposed | C1 | C1 | C1 | C1 | C1 | C1 |
| Initial particle images (no.) | 1,852,732 | 1,852,732 | 2,570,871 | 2,570,871 | 1,745,519 | 1,745,519 |
| Final particle images (no.) | 19,464 | 107,150 | 232,210 | 33,353 | 332,747 | 65,608 |
| Map resolution (Å) | 3.9 | 3.6 | 3.0 | 3.5 | 3.7 | 4.2 |
| FSC threshold | 0.143 | 0.143 | 0.143 | 0.143 | 0.143 | 0.143 |
| Map resolution range (Å) | 47.27 to 2.17 | 28.34 to 2.04 | 27.6 to 1.71 | 44.5 to 2.13 | 10.21 to 2.36 | 13.3 to 2.7 |
| **Refinement** | | | | | | |
| Initial model used (PDB code) | 8ZP1<br>AlphaFold model | 8ZP2<br>AlphaFold model | 9XBY<br>AlphaFold model | 9XFO<br>AlphaFold model | 9XBY<br>AlphaFold model | 24GL<br>AlphaFold model |
| Model composition | | | | | | |
| Non-hydrogen atoms | 14,953 | 14,923 | 16,233 | 16,070 | 16,027 | 16,052 |
| Protein residues | 1,917 | 1,908 | 2,081 | 2,061 | 2,055 | 2,059 |
| Ligands | 0 | 0 | 0 | 0 | 0 | |
| *B* factors (Å$^2$) | | | | | | |
| Protein | 219.73 | 221.41 | 149.34 | 200.07 | 214.96 | 279.27 |
| Ligand | 0 | 0 | 0 | 0 | 0 | 0 |
| R.m.s. deviations | | | | | | |
| Bond lengths (Å) | 0.004 | 0.006 | 0.006 | 0.006 | 0.004 | 0.006 |
| Bond angles (°) | 0.732 | 0.771 | 1.121 | 1.080 | 0.720 | 1.176 |
| **Validation** | | | | | | |
| MolProbity score | 2.28 | 2.48 | 1.80 | 1.92 | 2.01 | 2.03 |
| Clashscore | 13.57 | 14.15 | 7.81 | 11.04 | 9.91 | 19.36 |
| Poor rotamer (%) | 2.37 | 3.61 | 1.49 | 1.86 | 2.32 | 0.23 |
| Ramachandran plot | | | | | | |
| Favored (%) | 94.79 | 94.17 | 96.37 | 97.12 | 96.66 | 96.34 |
| Allowed (%) | 5.21 | 5.83 | 3.54 | 2.78 | 3.29 | 3.47 |
| Disallowed (%) | 0 | 0 | 0.1 | 0.1 | 0.05 | 0.20 |

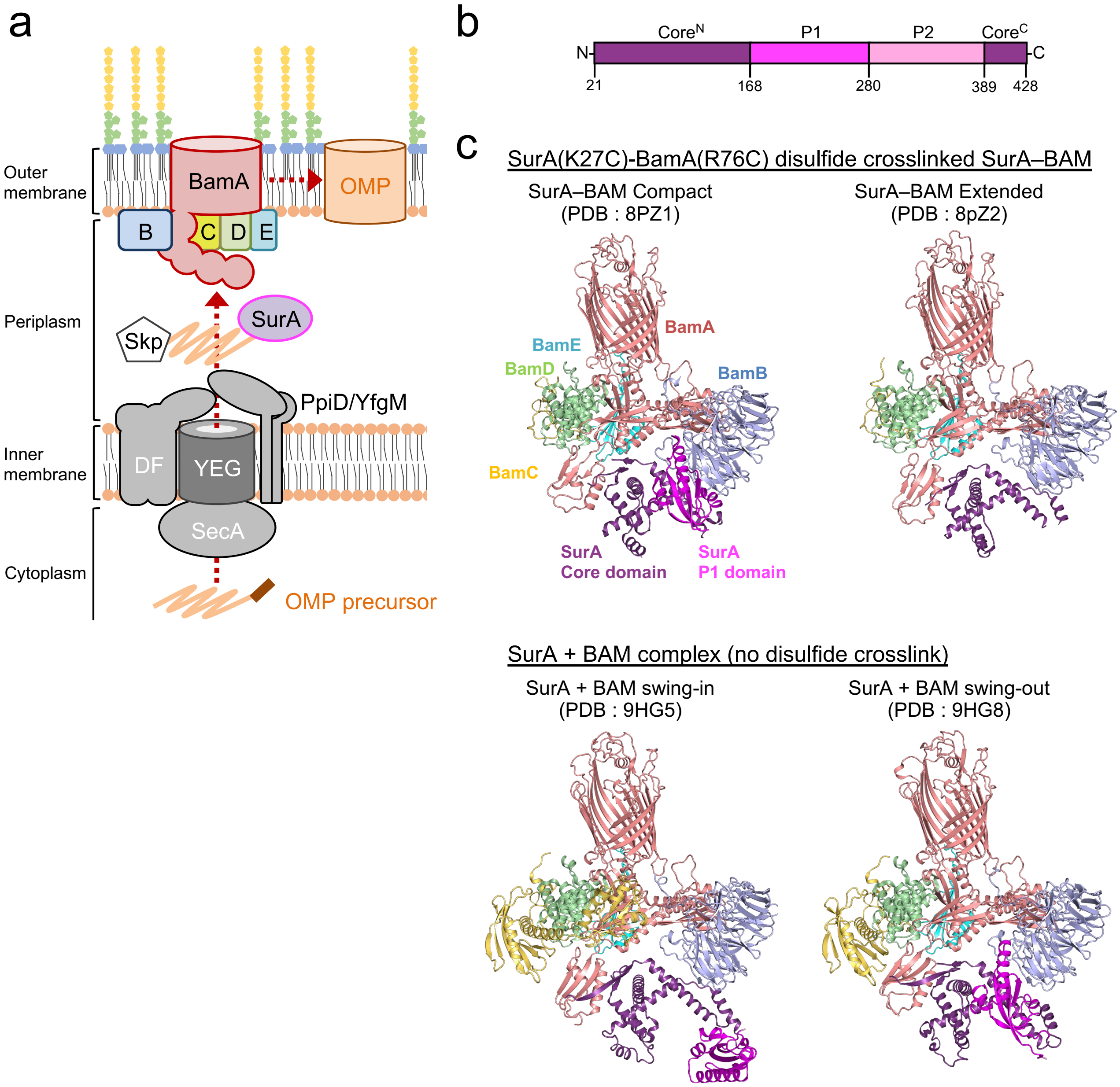


**Supplementary Fig. 1 | OMP assembly pathway and previous structures of the SurA–BAM complexes.**
**a** OMP assembly pathway. Precursor proteins synthesized in the cytoplasm with signal peptides are translocated via the SecYEG translocon. SecA, SecDF and Ppid/YfgM facilitate the protein translocation. In the periplasm, several chaperones, such as Skp and SurA, interact with translocated proteins and deliver them to the BAM complex. The BAM complex, consisting of BamA–E, mediates the insertion and assembly of OMPs into the outer membrane. **b** Schematic representations of three domains of SurA. **c** Previous SurA–BAM cryo-EM structures. SurA Core, and P1 domains are colored in purple and magenta, respectively. BamA, BamB, BamC, BamD, and BamE are in red, light blue, yellow, light green, and cyan, respectively. In all models, the P2 domain of SurA was not modeled, and in one of them (PDB ID: 8QZ2), the P1 domain was also not modeled.

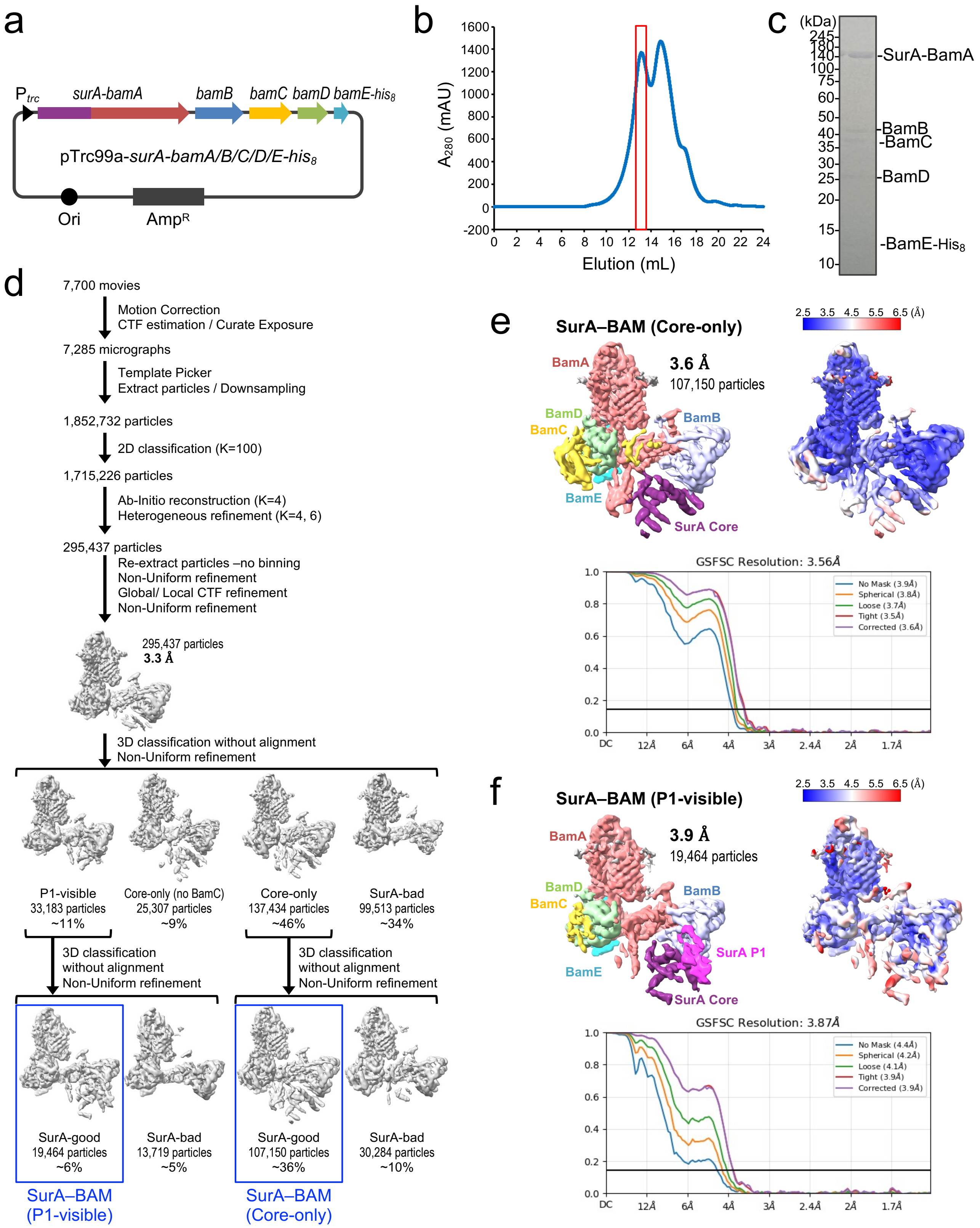


**Supplementary Fig. 2 | Cryo-EM analysis of the fused SurA–BAM.**
**a** Schematic of a plasmid encoding *surA-bamA/B/C/D/E-his*$_8$. **b** Size-exclusion chromatogram of the purified SurA–BAM complex using a Superose 6 Increase column. **c** SDS–PAGE analysis of the purified SurA–BAM complex stained with CBB. **d** Cryo-EM data-processing workflow for the SurA–BAM complex. **e, f** Final cryo-EM maps of Core-only and P1-visible SurA–BAM complexes. The maps are displayed at contour level 0.15 and 0.16, respectively. SurA Core, P1 and P2 domains are colored in purple, magenta and pink, respectively. BamA, BamB, BamC, BamD, and BamE are in red, light blue, yellow, light green, and cyan, respectively. The local resolution of the cryo-EM maps were calculated using CryoSPARC and are shown. Gold-standard Fourier shell correlation (GSFSC) curve used for global-resolution estimates within CryoSPARC are also shown.

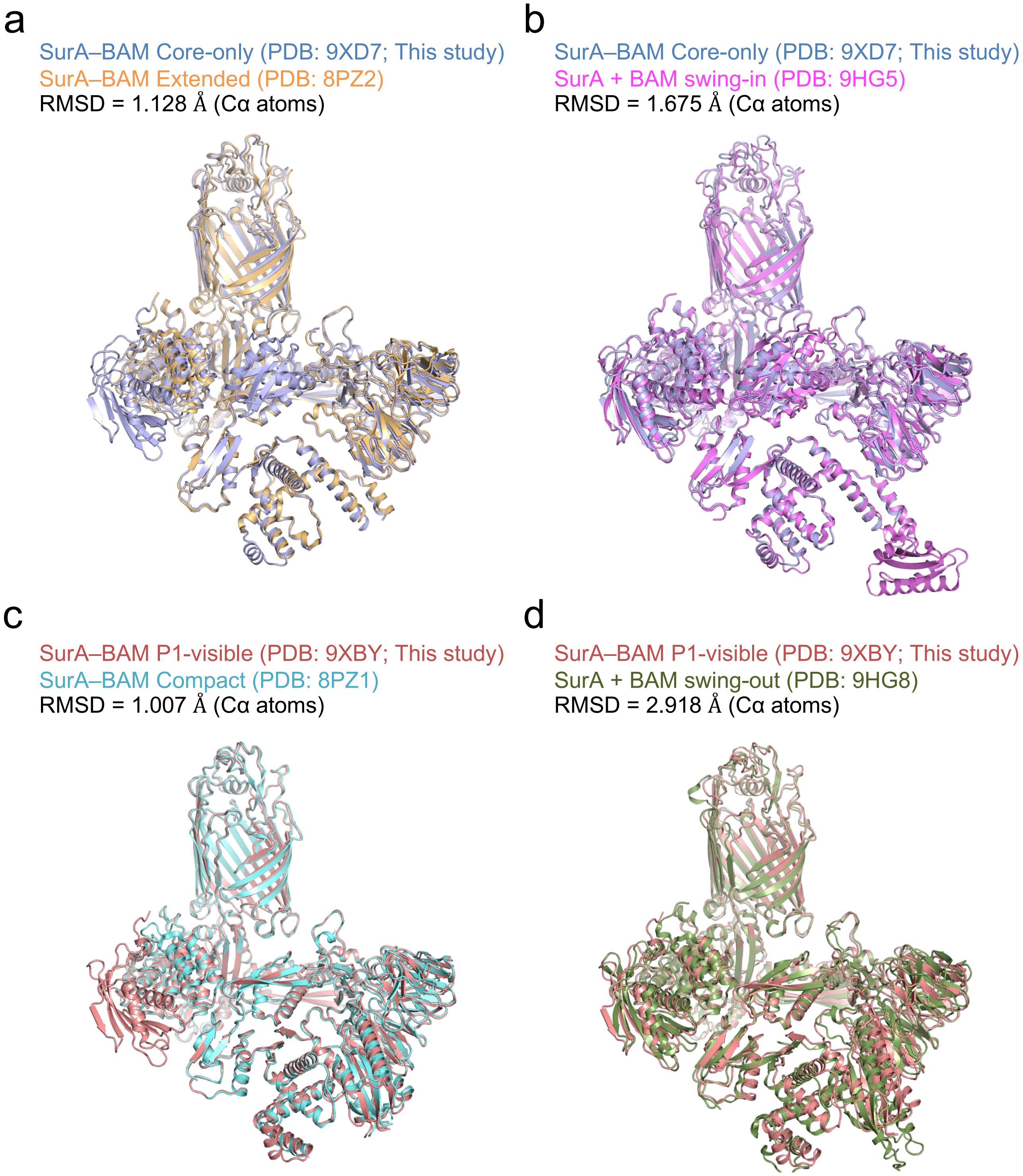


**Supplementary Fig. 3 | Structural comparisons of the SurA–BAM complexes.**
**a, b** A comparison of the Core-only SurA–BAM structure (PDB: 9XD7) with the extended structure (PDB: 8PZ2) (**a**) or with the swing-in structure (PDB: 9HG5) (**b**). **c, d** A comparison of the P1-visible SurA–BAM structure (PDB: 9XBY) with the compact structure (PDB: 8PZ1) (**c**) or with the swing-out structure (PDB: 9HG8) (**d**).

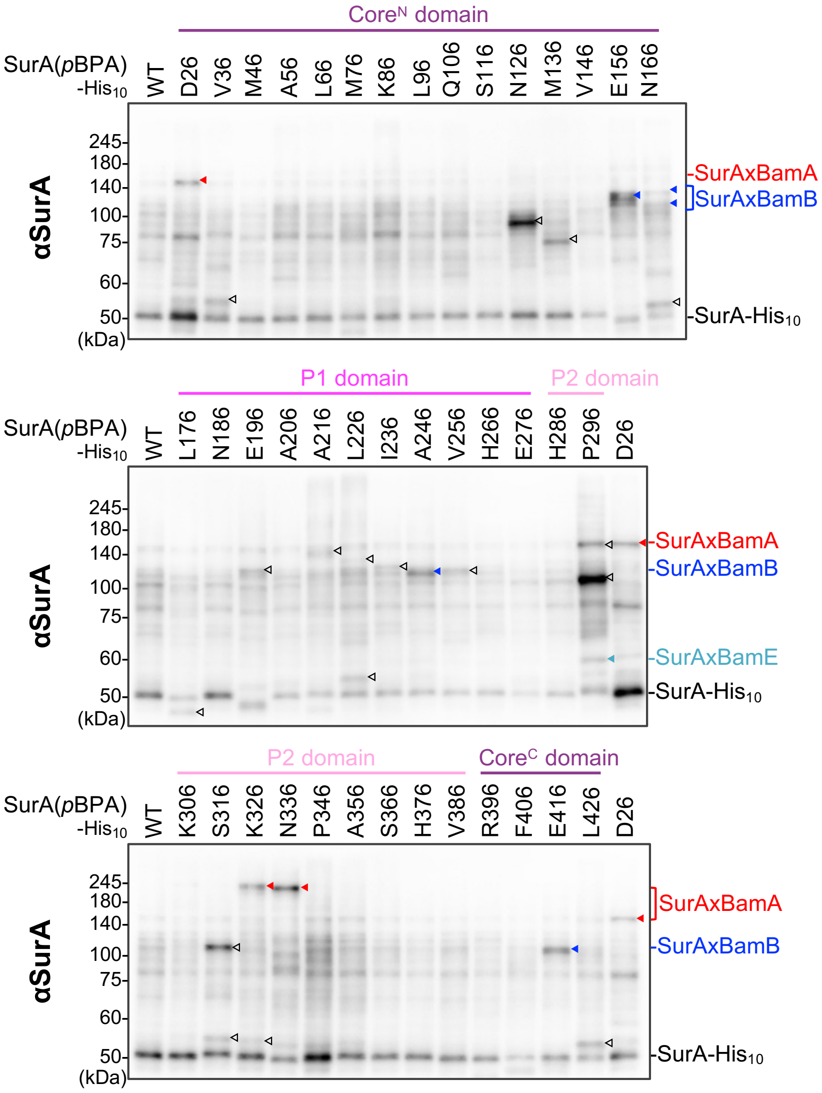


**Supplementary Fig. 4 | *In vivo* photo-crosslinking analysis of SurA.**
SN305 (Δ*surA*::*kan*) cells carrying both pEVOL-pBpF and pHM1021-*surA(amb)-his$_{10}$* were grown, induced with IPTG and UV-irradiated as described in Figs. 2b–d. Total cellular proteins were acid-precipitated and analyzed using SDS–PAGE followed by immunoblotting using anti-SurA antibodies. Red, blue, and cyan arrowheads indicate crosslinked products with BamA, BamB, and BamE, respectively. Open arrowheads indicate unidentified crosslinked products.

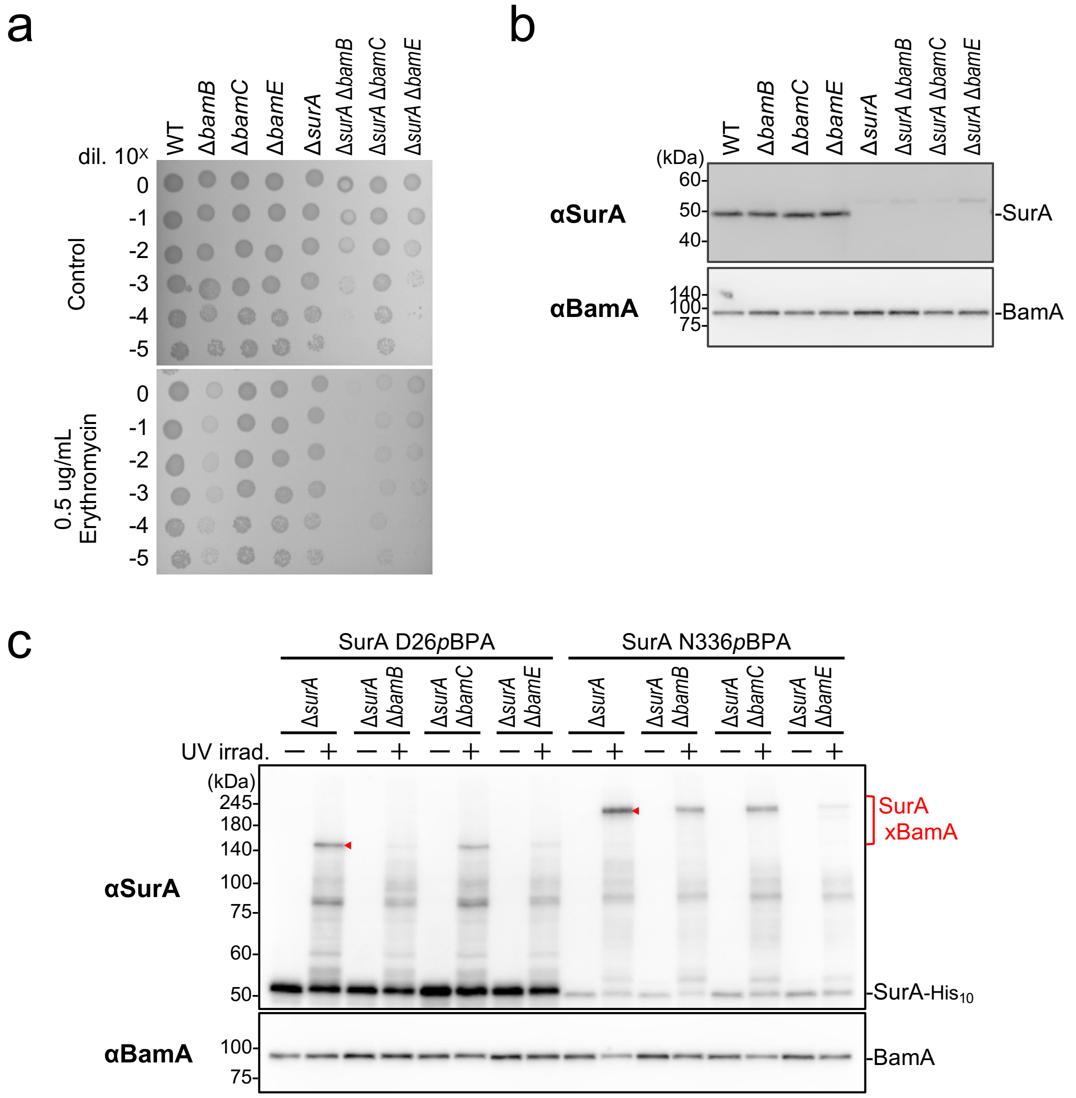


**Supplementary Fig. 5 | Functional relationship between SurA and Bam factors.**
**a** Synthetic phenotype of deletion mutations of *surA* and *bam* factors. Cells of AD16 derivatives were grown at 30 ºC in LB medium for 2.5 h. The cells were subsequently washed, suspended in saline, and serially diluted with saline (to approximately $10^9$ cells/mL). A total of 2.5 μL of each of the diluted cells was spotted on an LB agar plate with or without erythromycin. The plates were incubated at 30 ºC for 22 h. **b** Same cells in (**a**) were grown at 30 ºC in LB medium for 2.5 h. Total cellular proteins were acid-precipitated and analyzed by 10% Laemmli SDS–PAGE followed by immunoblotting using the indicated antibodies. **c** Effect of deletion mutations of Bam factors on SurA–BamA crosslinking. Cells of RM5457 (Δ*surA*) derivatives carrying both pEVOL-pBpF and pHM1021-*surA(amb)-his*$_{10}$ were grown, induced with IPTG, and UV-irradiated as described in Figs. 2b–d. Total cellular proteins were acid-precipitated and analyzed using SDS–PAGE followed by immunoblotting with the indicated antibodies.

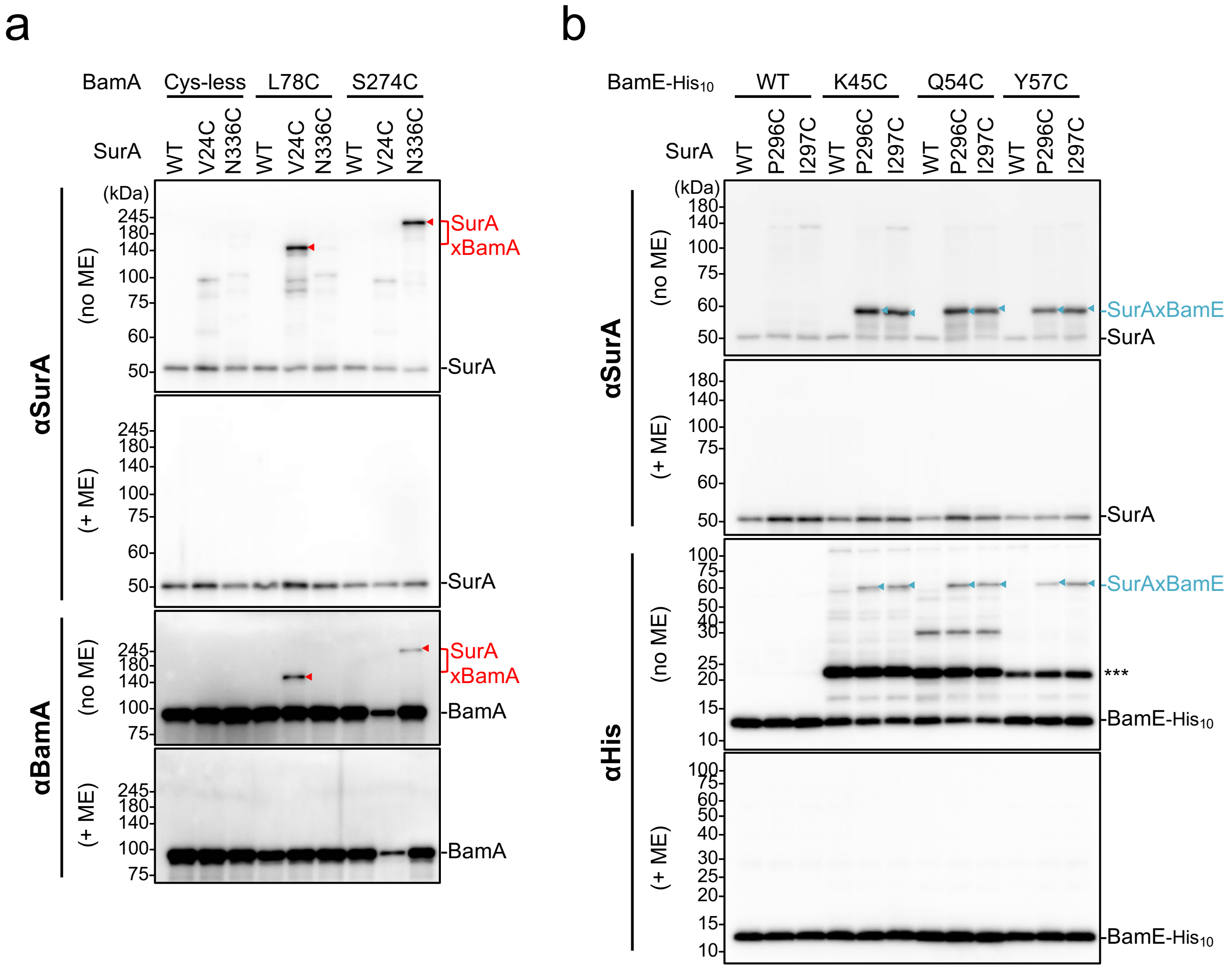


**Supplementary Fig. 6 | *In vivo* disulfide crosslinking between SurA and the BAM complex.**

**a** SurA–BamA disulfide crosslinking. RM4728 (*bamA* under arabinose promoter) cells carrying a combination of plasmids encoding WT or a Cys-introduced mutant of SurA and BamA as indicated were grown at 30 ºC in LB medium and induced with 1 mM IPTG for 3 h to express SurA(Cys) and BamA(Cys). Total cellular proteins were acid-precipitated, solubilized with SDS-buffer containing NEM (for blocking free thiol groups), treated with or without β-mercaptoethanol (ME), and analyzed by 7.5% Laemmli SDS–PAGE under reducing (+ ME) or nonreducing (no ME) conditions and immunoblotting with the indicated antibodies. **b** SurA–BamE disulfide crosslinking. RM5475 (Δ*surA*, Δ*bamE*) cells carrying a combination of plasmids encoding WT or a Cys-introduced mutant of SurA and BamE-$His_{10}$ as indicated were grown at 30 ºC in LB medium and induced with 1 mM IPTG for 2.5 h to express SurA(Cys) and BamE(Cys)-$His_{10}$. Total cellular proteins were acid-precipitated and analyzed as described in (**a**).

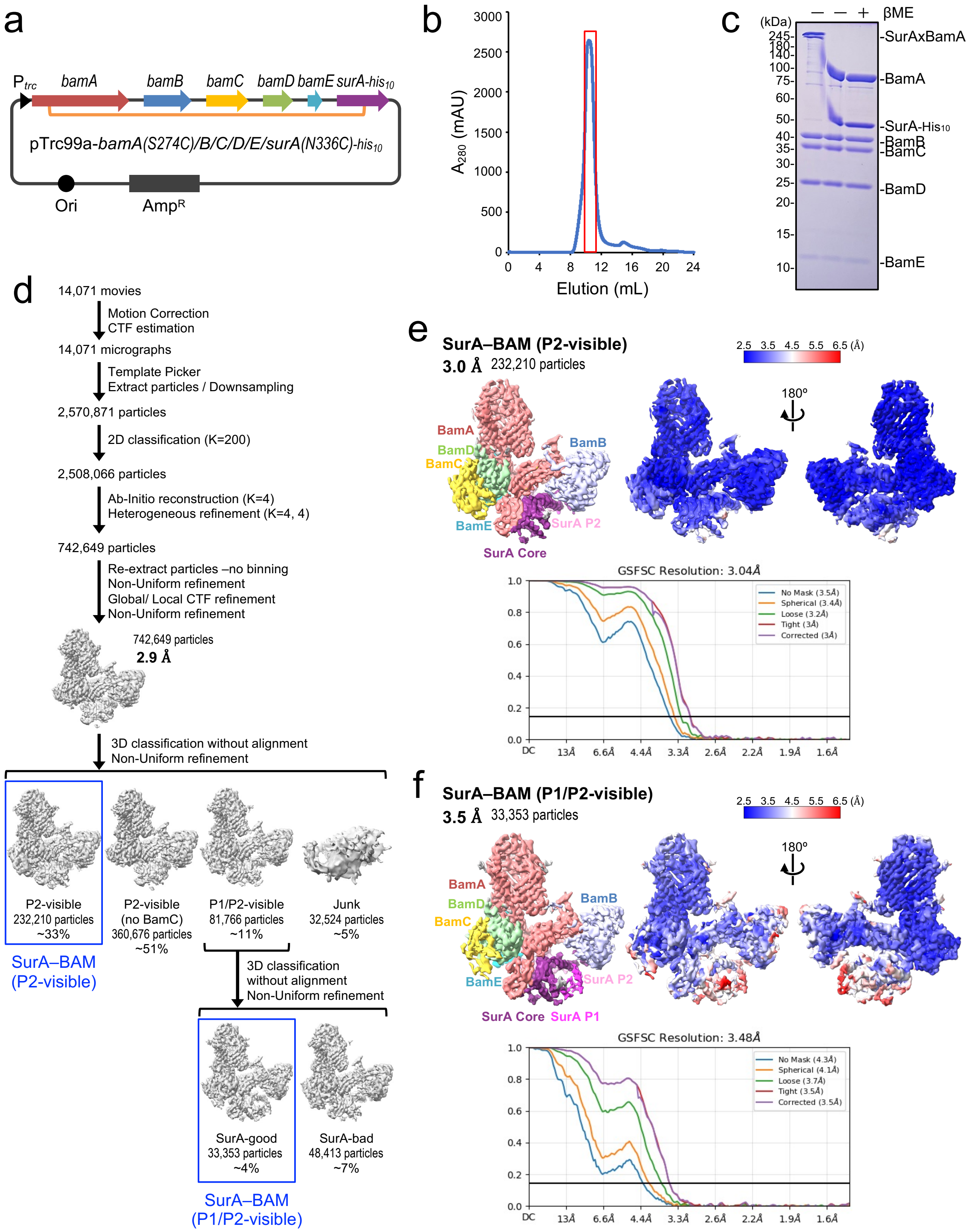


**Supplementary Fig. 7 | Cryo-EM analysis of the disulfide crosslinked SurA–BAM complex 1.**
**a** Schematic of a plasmid encoding *bamA(S274C)/B/C/D/E/surA(N336C)-his$_{10}$*. **b** Size-exclusion chromatogram of the purified SurA–BAM complex using a Superdex 200 Increase column. **c** SDS–PAGE analysis of the purified SurA–BAM complex stained with CBB. ME was added to the rightmost lane. **d** Cryo-EM data-processing workflow for the SurA–BAM complex. **e, f** Final cryo-EM maps of P2-visible and P1/P2-visible SurA–BAM complexes. The maps are displayed at contour level 0.08. SurA Core, P1 and P2 domains are colored in purple, magenta and pink, respectively. BamA, BamB, BamC, BamD, and BamE are in red, light blue, yellow, light green, and cyan, respectively. The local resolution of the cryo-EM maps were calculated using CryoSPARC and are shown. Gold-standard Fourier shell correlation (GSFSC) curve used for global-resolution estimates within CryoSPARC are also shown.

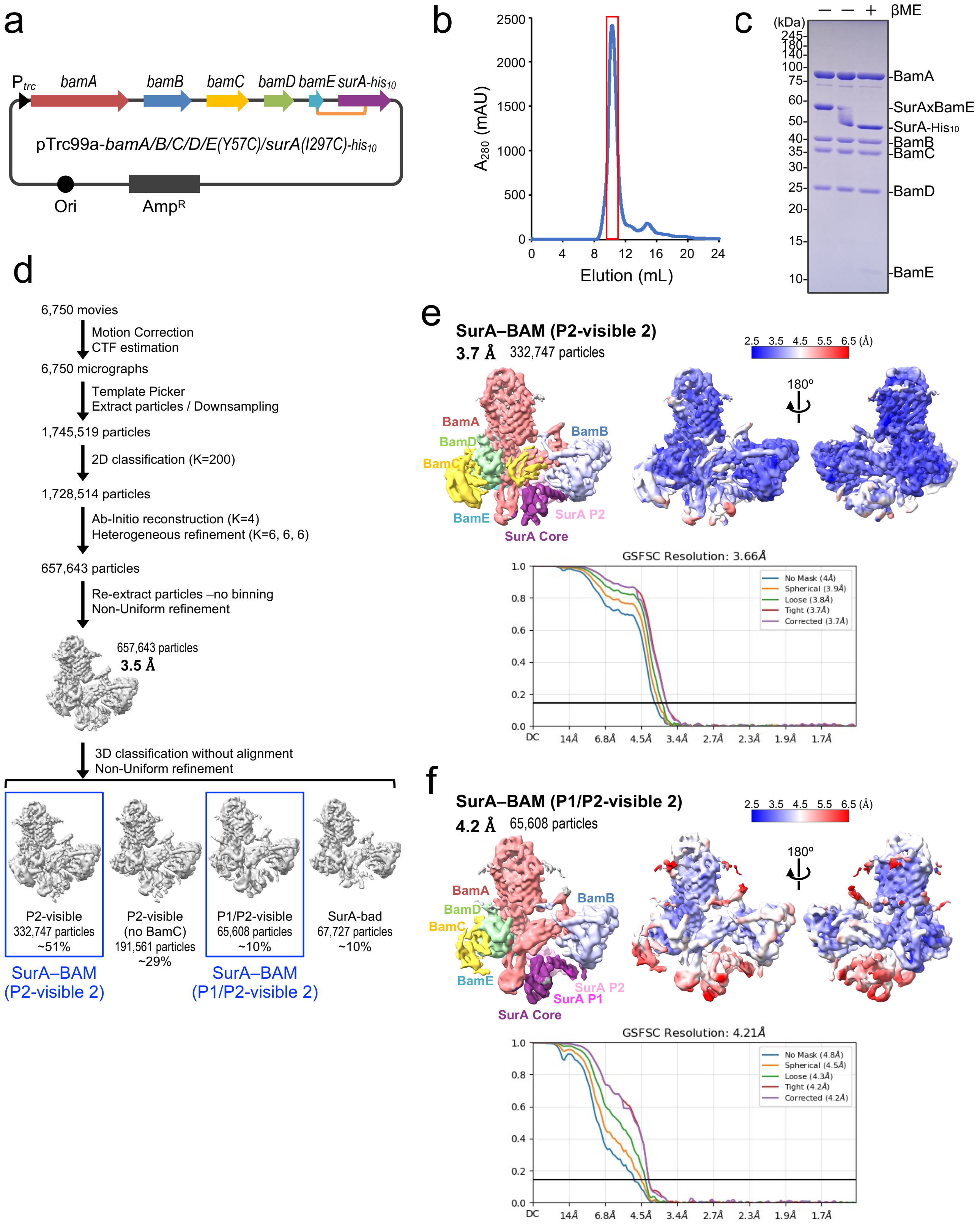


**Supplementary Fig. 8 | Cryo-EM analysis of the disulfide crosslinked SurA–BAM complex 2.**
**a** Schematic of a plasmid encoding *bamA/B/C/D/E(Y57C)/surA(I297C)-his*$_{10}$. **b** Size-exclusion chromatogram of the purified SurA–BAM complex using a Superdex 200 Increase column. **c** SDS–PAGE analysis of the purified SurA–BAM complex stained with CBB. ME was added to the rightmost lane. **d** Cryo-EM data-processing workflow for the SurA–BAM complex. **e, f** Final cryo-EM maps of P2-visible 2 and P1/P2-visible 2 SurA–BAM complexes. The maps are displayed at contour level 0.055. SurA Core, P1 and P2 domains are colored in purple, magenta and pink, respectively. BamA, BamB, BamC, BamD, and BamE are in red, light blue, yellow, light green, and cyan, respectively. The local resolution of the cryo-EM maps were calculated using CryoSPARC and are shown. Gold-standard Fourier shell correlation (GSFSC) curve used for global-resolution estimates within CryoSPARC are also shown.

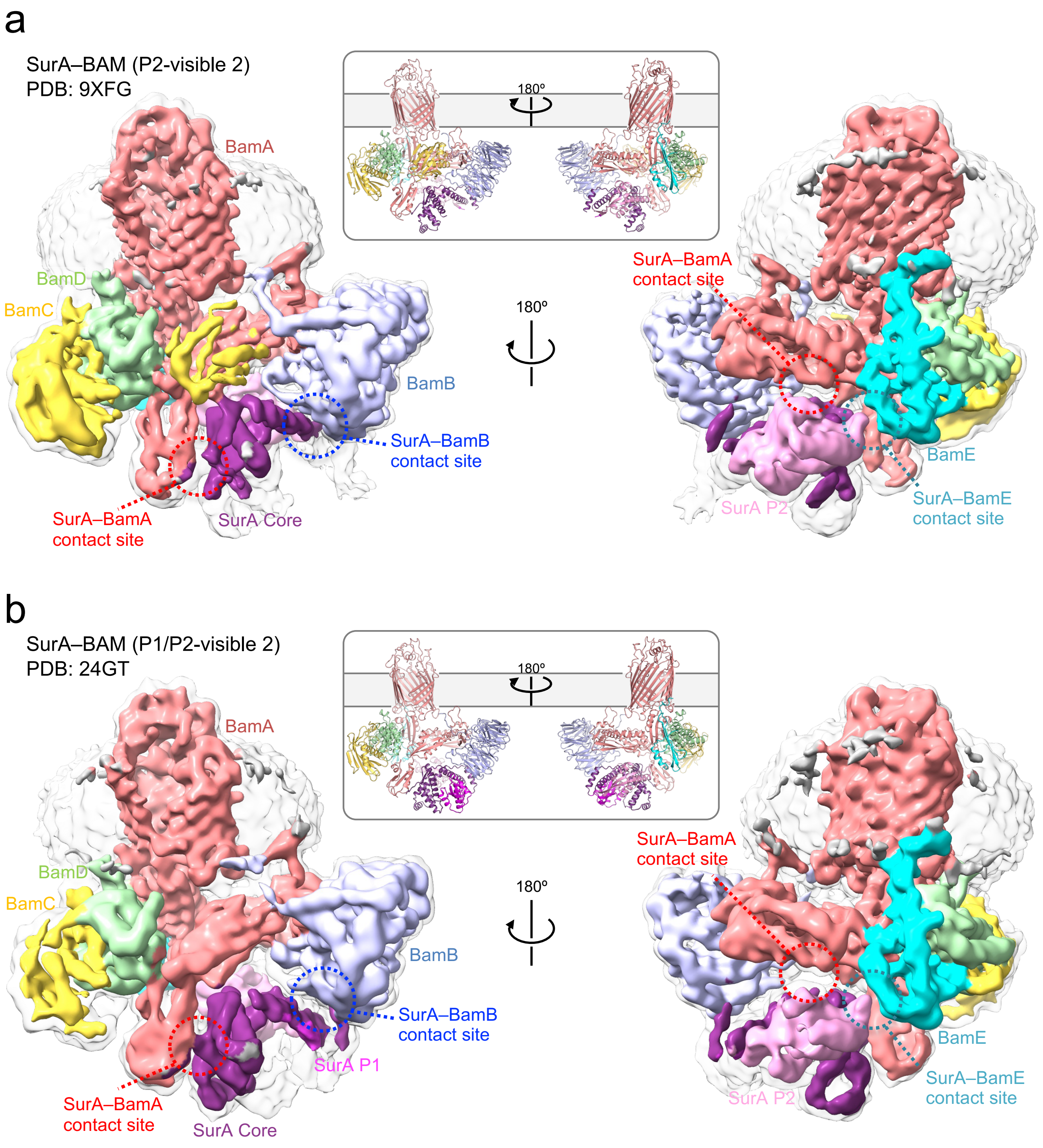


**Supplementary Fig. 9 | Cryo-EM structures of the P2-visible 2 and P1/P2-visible 2 SurA–BAM complexes.**
**a** Cryo-EM map and cartoon model of the SurA–BAM P2-visible 2 complex (PDB: 9XFG). **b** Cryo-EM map and cartoon model of the SurA–BAM P1/P2-visible complex (PDB: 24GT). The SurA Core, P1 and P2 domains are colored in purple, magenta and light pink, respectively. BamA, BamB, BamC, BamD, and BamE are colored as in Fig. 1. Contact sites of SurA with BamA, BamB and BamE are shown in red, blue and cyan circles, respectively.

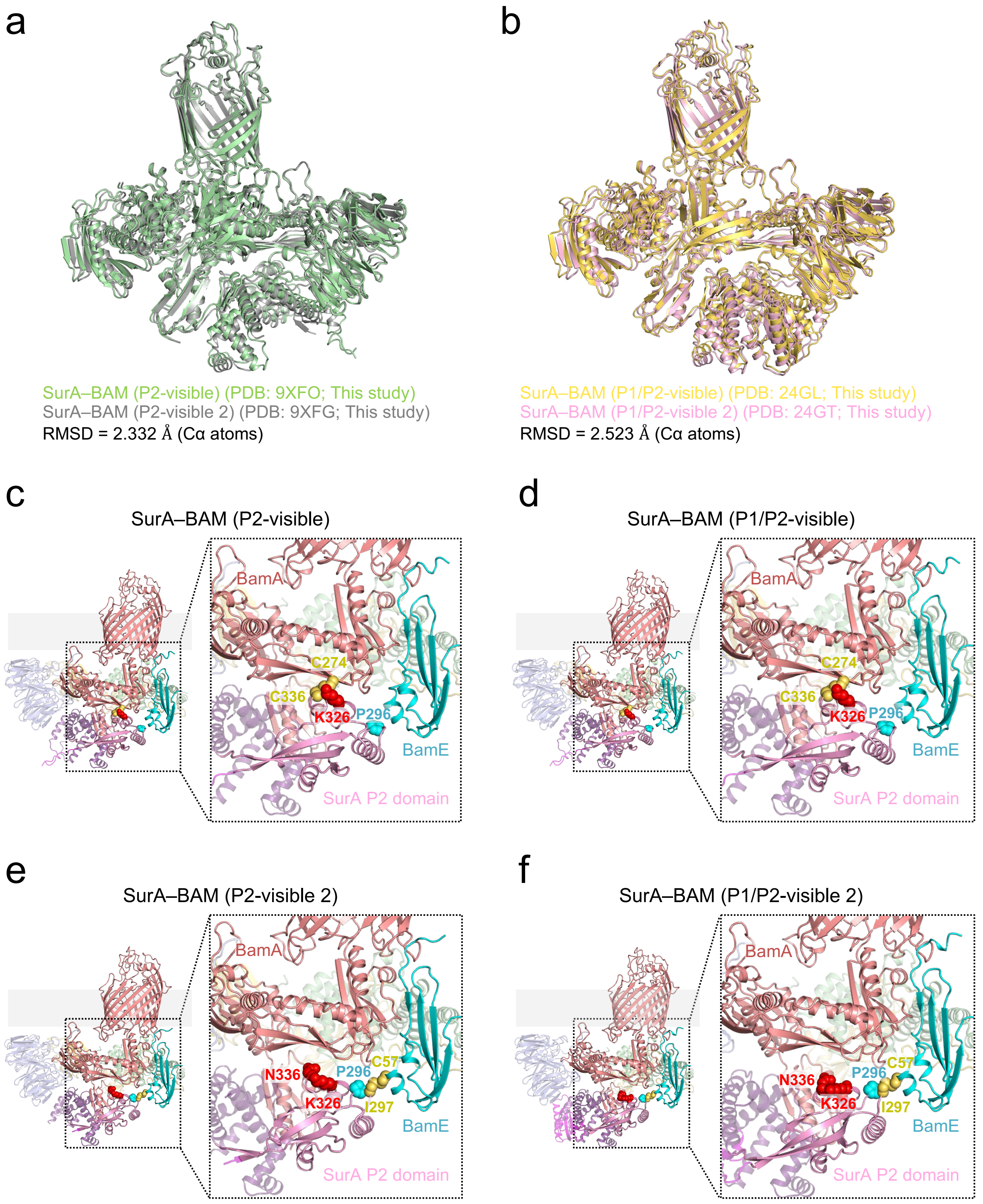


**Supplementary Fig. 10 | Structural details of cryo-EM structures of the P2-visible and P1/P2-visible SurA/BAM complexes.**

**a,b** Comparison of the cryo-EM structures of the P2-visible or P1/P2-visible SurA–BAM complexes. **c-f** Mapping of the crosslinking sites onto the SurA–BAM cryo-EM structures. SurA residues crosslinked with BamA and BamE are indicated by red and cyan spheres, respectively. Disulfide-bonded residues are colored in yellow spheres.

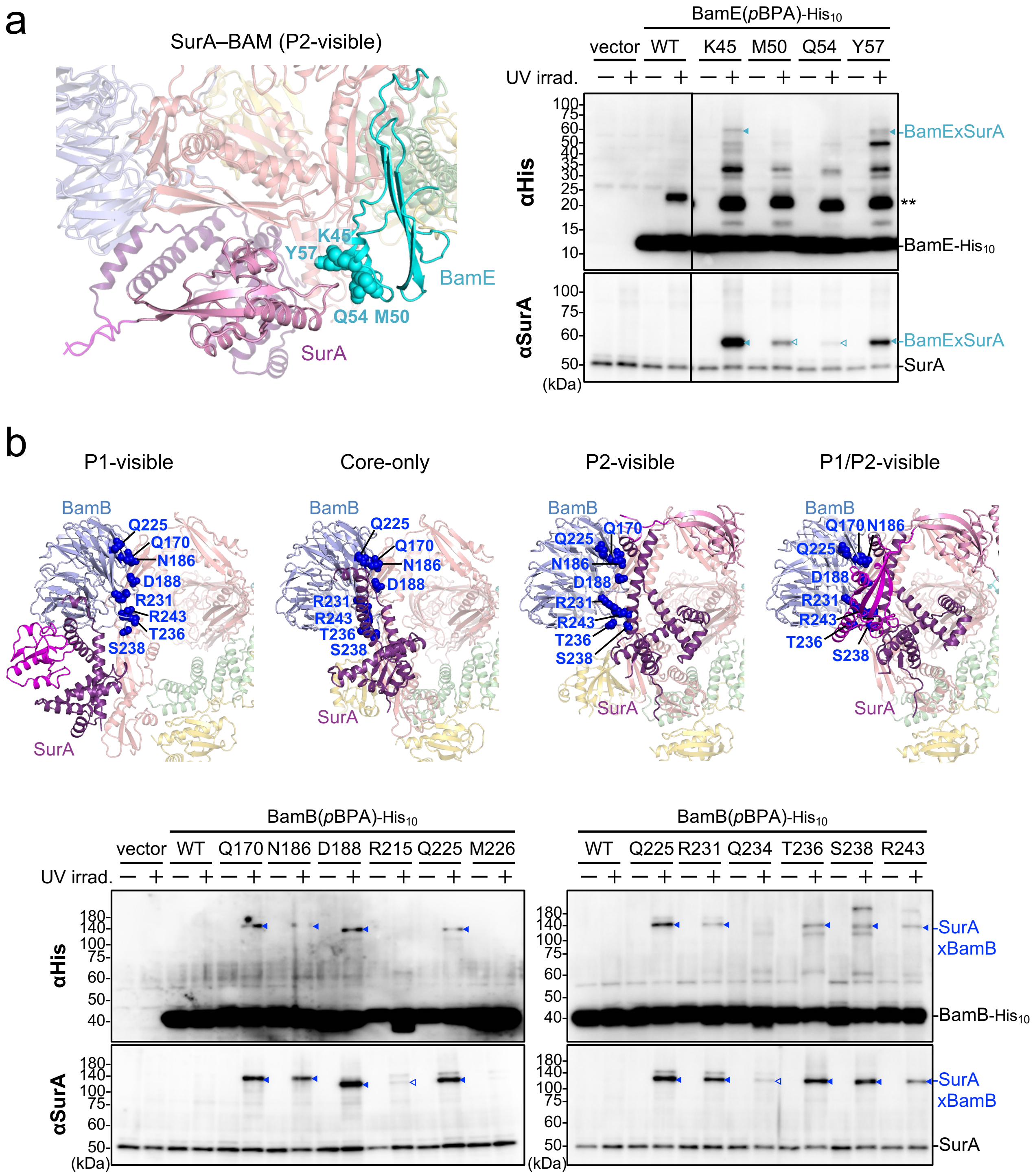


**Supplementary Fig. 11 | *In vivo* photo-crosslinking between SurA and BamE or BamB.**

**a** *In vivo* photo-crosslinking analysis of BamE(*p*BPA) variants. (left) Mapping of the *p*BPA-incorporated sites of BamE on the SurA–BAM P2-visible cryo-EM structure (PDB: 9XFO). (right) SN535 (Δ*bamE*) cells carrying both pEVOL-pBpF and pHM1021 (vector) or pHM1021-*bamE(amb)-$his_{10}$* were grown at 30 ºC in LB medium containing 0.02% arabinose and 0.5 mM *p*BPA until early log phase and induced with 1 mM IPTG for 1 h to express the indicated BamB(*p*BPA) variants. Cultures were then divided into two portions and treated with or without UV irradiation for 10 min at 4 ºC. Total cellular proteins were acid-precipitated and analyzed using SDS–PAGE followed by immunoblotting using indicated antibodies. ** indicates a nonspecific band. **b** *In vivo* photo-crosslinking analysis of BamB(*p*BPA) variants. (upper) Mapping of the *p*BPA-incorporated sites of BamB on the SurA–BAM cryo-EM structures. (lower) SN147 (Δ*bamB*) cells carrying both pEVOL-pBpF and pHM1021 (vector) or pHM1021-*bamB(amb)-$his_{10}$* were grown and analyzed using SDS-PAGE followed by immunoblotting using the indicated antibodies as described in (**a**).

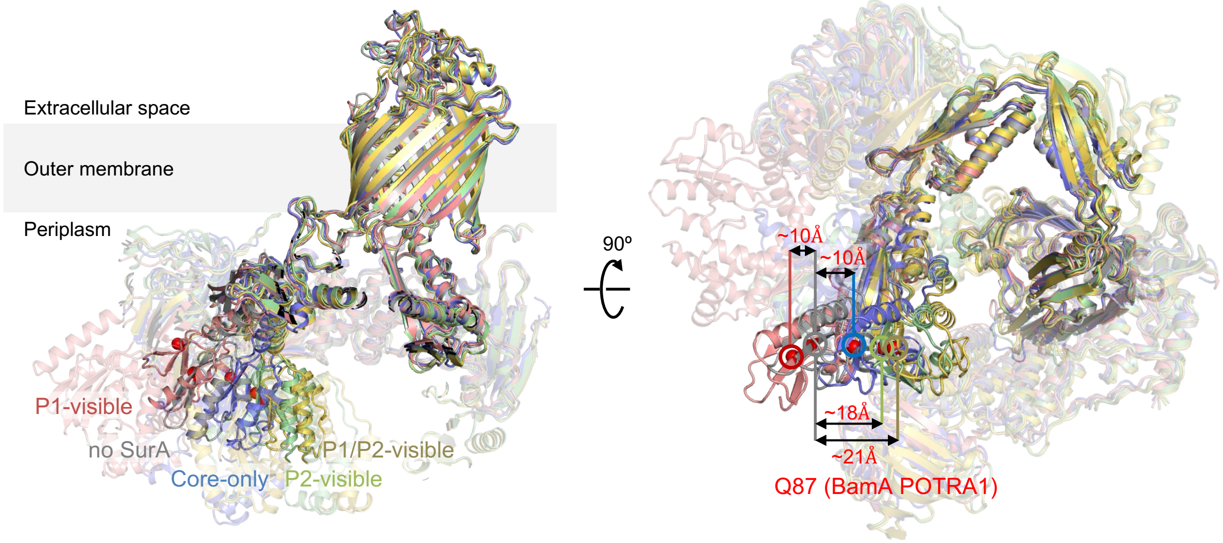


**Supplementary Fig. 12 | Comparison of BamA in our SurA–BAM cryo-EM structures with a SurA-lacking BAM structure.**

Cryo-EM structures of the SurA–BAM complex are superposed: P1-visible (red, PDB: 9XBY), Core-only (light blue, PDB: 9XD7), P2-visible (light green, PDB: 9XFO), P1/P2-visible (yellow, PDB: 24GL), and SurA-lacking BAM (gray, PDB: 9CNW). Distances between the Q87 Cα position in the SurA-lacking BAM structure and the corresponding Q87 Cα positions in the BamA POTRA1 domain of each SurA–BAM structure are indicated.

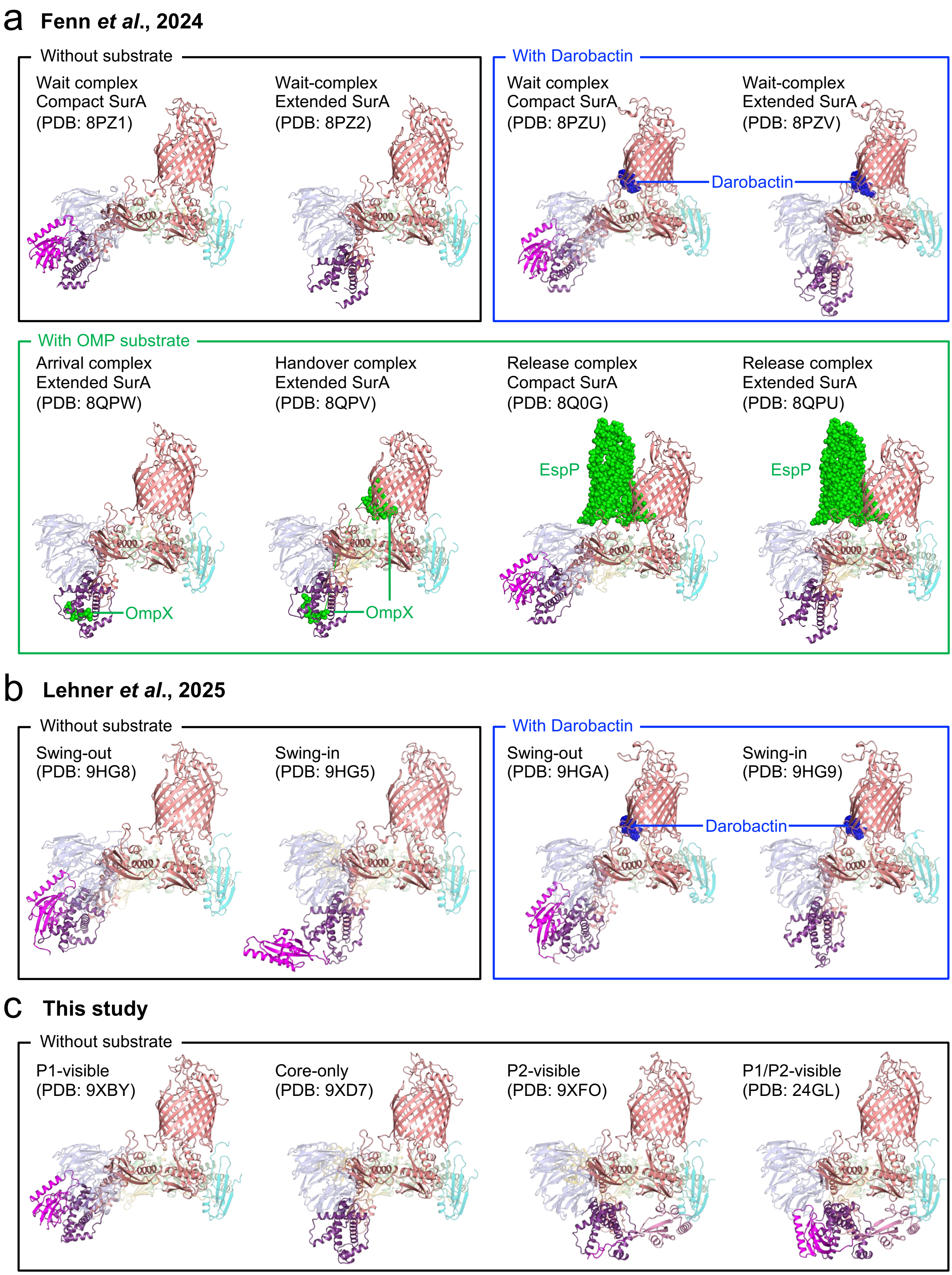


**Supplementary Fig. 13 | Cryo-EM structures of the SurA–BAM complex.**
**a** SurA-BAM cryo-EM structures previously reported in Fenn *et al.*, 2024[16]. **b** SurA-BAM cryo-EM structures previously reported in Lehner *et al.*, 2025[17]. **c** SurA-BAM cryo-EM structures determined in this study. The SurA-BAM complexes are shown as cartoon models. The SurA Core, P1 and P2 domains are colored in purple, magenta and light pink, respectively. BamA, BamB, BamC, BamD, and BamE are colored in red, light blue, yellow, light green and cyan, respectively. Substrate OMPs and darobactin, a BAM inhibitor, are shown in green and blue spheres, respectively.

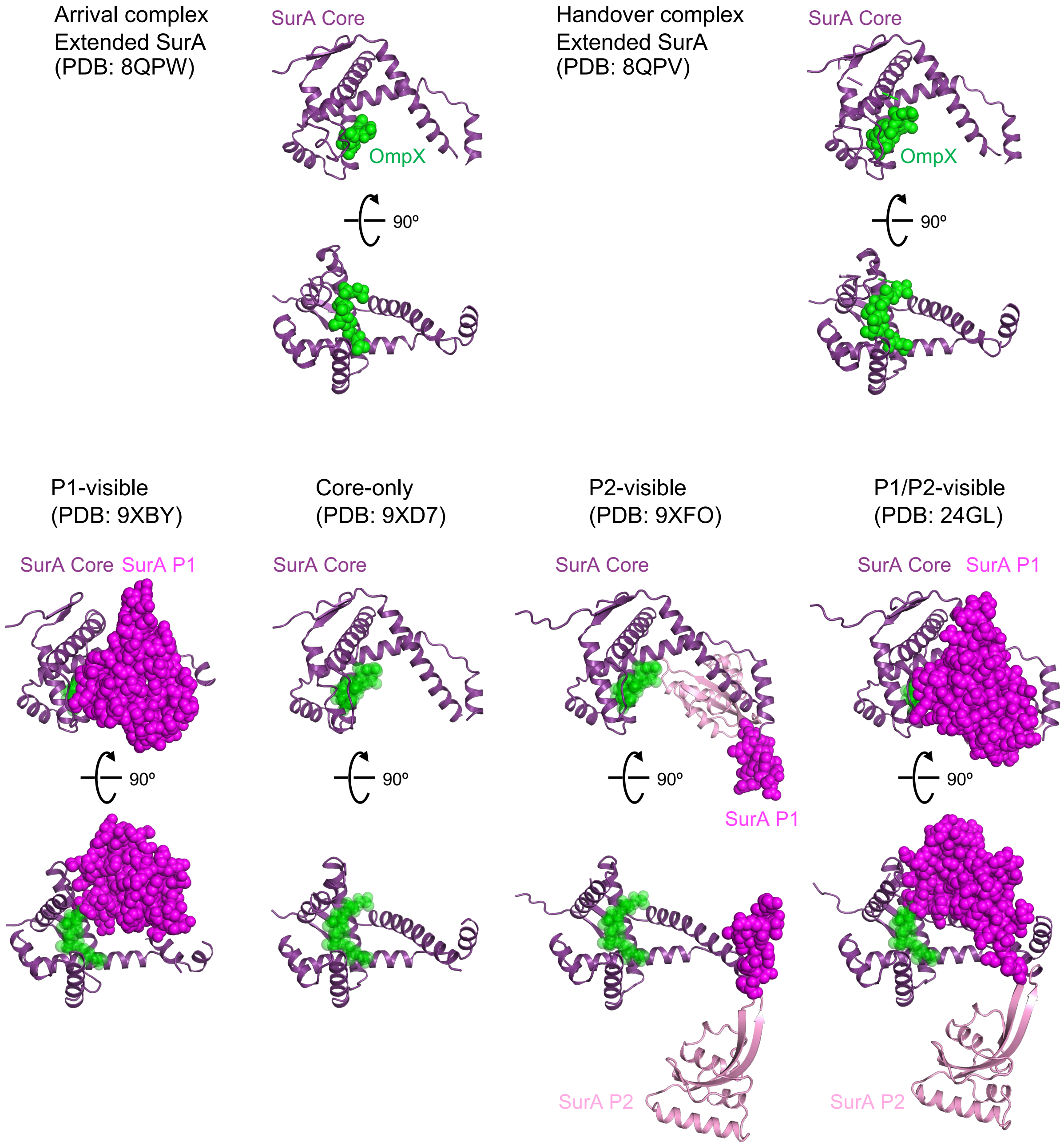


**Supplementary Fig. 14 | Substrate binding to the SurA Core domain in SurA–BAM cryo-EM structures.**
The SurA core domain and a substrate polypeptide in previously-reported substrate-bound SurA-BAM complexes (PDB: 8QPW and 8QPV; upper panels) are compared with those in our substrate-free SurA–BAM complexes. The SurA Core and P2 domains are shown as cartoon models and colored in purple and light pink, respectively. Substrate OMP polypeptides and the SurA P1 domain are shown in green and magenta spheres, respectively. OmpX polypeptides from Handover complex are superimposed onto our SurA-BAM structures are shown as ghost models with 50% transparency (lower panels). In P1-visible and P1/P2-visible structures, the OmpX polypeptide and the P1 domain show a steric hindrance.
